# Combined use of metagenomic sequencing and host response profiling for the diagnosis of suspected sepsis

**DOI:** 10.1101/854182

**Authors:** Henry K. Cheng, Susanna K. Tan, Timothy E. Sweeney, Pratheepa Jeganathan, Thomas Briese, Veda Khadka, Fiona Strouts, Simone Thair, Sudeb Dalai, Matthew Hitchcock, Ashrit Multani, Jenny Aronson, Tessa Andermann, Alexander Yu, Samuel Yang, Susan P. Holmes, W. Ian Lipkin, Purvesh Khatri, David A. Relman

## Abstract

**Background:** Current diagnostic techniques are inadequate for rapid microbial diagnosis and optimal management of patients with suspected sepsis. We assessed the clinical impact of three powerful molecular diagnostic methods.

**Methods:** With blood samples from 200 consecutive patients with suspected sepsis, we evaluated 1) metagenomic shotgun sequencing together with a Bayesian inference approach for contaminant sequence removal, for detecting bacterial DNA; 2) viral capture sequencing; and 3) transcript-based host response profiling for classifying patients as infected or not, and if infected, with bacteria or viruses. We then evaluated changes in diagnostic decision-making among three expert physicians by unblinding the results of these methods in a staged fashion.

**Results:** Metagenomic shotgun sequencing confirmed positive blood culture results in 14 of 26 patients. In 17 of 200 patients, metagenomic sequencing and viral capture sequencing revealed organisms that were 1) not detected by conventional hospital tests within 5 days after presentation, and 2) classified as of probable clinical relevance by physician consensus. Host response profiling led at least two of three physicians to change their diagnostic decisions in 46 of 100 patients. The data suggested possible bacterial DNA translocation in 8 patients who were originally classified by physicians as noninfected and illustrate how host response profiling can guide interpretation of metagenomic shotgun sequencing results.

**Conclusions:** The integration of host response profiling, metagenomic shotgun sequencing, and viral capture sequencing enhances the utility of each, and may improve the diagnosis and management of patients with suspected sepsis.

---

The early recognition and diagnosis of severe infection and sepsis is a significant clinical priority. Despite advances in microbial detection methods, clinicians typically rely on presumptive clinical diagnoses and empiric therapy with broad-spectrum antimicrobials, increasing the risks for adverse drug effects^1^ and the development of antimicrobial resistance. Two emerging approaches, metagenomic sequencing and host response profiling, may each promote the rapid diagnosis of sepsis. Their use in a prospective fashion, and especially in combination, has not been adequately assessed and deserves careful study.

In theory, metagenomic sequencing can identify any microorganism to the species- or strain-level without the need for a prior hypothesis or reliance on cultivation, as long as there are nucleic acids of sufficient abundance and length from the organism(s) in the specimen. Case reports, validation, and interventional studies have highlighted the potential power of this approach^2–7^. Some methods incorporate microbial enrichment or human depletion steps in order to improve ‘signal to noise’ ratios^8,9^. For example, viral capture sequencing for vertebrate viruses (VirCapSeq-VERT) is a metagenomic sequencing approach that enriches for all 207 viral taxa known to infect vertebrates (including humans) with sensitivity similar to the real-time polymerase chain reaction assays currently employed in clinical microbiology laboratories^10,11^.

The mere presence of specific molecular components of an infectious agent in a patient is insufficient however to incriminate the agent as the cause of that patient’s disease^12,13^. For example, the presence of bacterial nucleic acids in a specimen of blood could be explained by contamination of the specimen with skin bacteria or their DNA during collection^14^, or even normal low-level translocation of commensal bacteria or their components into the bloodstream during states of health^15^. Viral sequences may represent latent or clinically-irrelevant viruses in circulating blood cells or their nucleic acids in plasma. Contamination of specimens with microbial nucleic acids from laboratory reagents at the time of specimen processing has been shown to critically affect results in the study of low-microbial biomass samples, such as blood^16^. Finally, false-positive and -negative results may reflect bioinformatic errors^17^ and faulty reference databases^3,18^, or other technical errors. The failure to address these same challenges in the use of other nucleic acid-based testing approaches such as multiplex pathogen PCR panels and *C. difficile* PCR testing has led to unnecessary antimicrobial treatments, delayed diagnoses, and/or detrimental patient outcomes^19–22^. The risks of these adverse outcomes are magnified with metagenomic approaches because of their broad range and the ubiquity of microbial nucleic acids.

Host RNA transcript-based profiles can provide evidence of a clinically relevant response to all infections or to broad classes of infectious agents, even though they may not be agent-specific.^23–27^ For this reason, host response profiling methods offer complementary benefits to methods for detecting microbes or their components^28^. Furthermore, host RNA signatures that distinguish infected from noninfected patients, and bacterial from viral infections^25,29,30^ are available to guide initial treatment, since relevant RNAs are expressed early in infection and can be measured rapidly.

We hypothesized that metagenomic sequencing and host response profiling would provide clinically useful information about the potential cause of suspected sepsis, that current, routine diagnostic tests fail to provide, and that their use in combination could prove complementary. Langelier et al. provided the first integration of these two approaches to diagnose lower respiratory tract infections.^31^ In our study, we prospectively enrolled 200 consecutive adult patients who presented to the Emergency Department with suspected sepsis^32^, and applied three molecular approaches to blood specimens: 1) metagenomic shotgun next-generation sequencing (mNGS) for detection of bacteria; 2) VirCapSeq-VERT for detection of DNA and RNA viruses; and 3) a previously-defined human response-based transcript signature^25^ for classifying patient infection status. We developed a Bayesian method for distinguishing blood- from reagent-associated DNA sequences in the mNGS data. Three infectious diseases physicians performed chart reviews on all patients in a blinded manner and then were provided results from the three diagnostic methods in a staged fashion. We report on the added value of these methods alone and together in generating clinically relevant diagnoses.

## METHODS

### Subject Enrollment

This study was approved by the Stanford University Administrative Panel on Human Subjects Research. Plasma and whole blood samples were collected prospectively from consecutive adult patients presenting to the Stanford University Hospital Emergency Department (ED) who were not pregnant; met 2 of 4 SIRS criteria^32^; and were suspected to have infection by triage nurses or other clinicians. We then identified 200 patients (spanning 128 days in 2016) who met additional criteria (see Supplementary Text). We collected 2.5 ml of whole blood from each of 10 healthy adults to serve as controls for host response profiling.

### Metagenomic sequencing

DNA was extracted from 200-400 μL of plasma with the QIAamp Circulating Nucleic Acid Kit (QIAGEN). DNA extraction was performed in batches of 24, with 3-4 negative controls per batch, consisting of molecular-grade water, to monitor environmental and reagent contamination during sample processing. Libraries were prepared with the KAPA HyperPrep Kit (Roche), and sequenced on the HiSeq 4000 (Illumina) with 2×150 nucleotide paired-end reads.

In a pilot experiment, we sequenced DNA from a plasma sample from each of 15 patients with a positive bacterial blood culture, at a depth of 40M-60M reads/sample, and 4 negative controls at a depth of 2-6M reads/sample. We then sequenced plasma samples from the other 185 patients each to a depth of 10M-52M reads, alongside 36 negative control samples sequenced to a depth of 3M-6M reads.

Bacterial reads were classified at the species-level with Kraken^33^ using a conservative alignment threshold and were further analyzed with phyloseq^34^. Exploratory analysis using principal components analysis (PCA) showed possible batch-effects (Figs. S1, S2). To distinguish blood-associated DNA sequences from contaminant sequences, we developed a Bayesian statistical method that leverages data from negative control samples (Fig. S3). Further details on sample preparation, bioinformatics, exploratory analysis, and the contaminant identification algorithm are provided in Supplementary Text.

### VirCapSeq-VERT

Nucleic acid was extracted from 150 µl of plasma. VirCapSeq-VERT enriched libraries were sequenced on a HiSeq 2500, generating 1 × 100 nucleotide single end reads. Additional details on VirCapSeq-VERT sequencing and associated bioinformatics are available in Supplementary Text.

### Host RNA transcript profiling

We tested samples from 193 patients and 10 healthy adult volunteers with a previously described 18-gene host-response assay consisting of 1) an 11-gene set to distinguish noninfection- and infection-associated SIRS, the Sepsis MetaScore (SMS)^24^; and 2) a 7-gene set to distinguish bacterial and viral infections, the ‘bacterial-viral metascore’ (BVS)^25^. qRT-PCR was performed in triplicate using commercial TaqMan assays on the Biomark HD platform (Fluidigm). Samples from seven patients were not profiled because of PCR failure. SMS and bacterial-viral scores were calculated as previously described^25^. Since this was the first use of qRT-PCR for this 18-gene assay, SMS and BVS cutoffs were re-established with the data from this study (Fig. S4, S5). With these score cutoffs, host response classifications of ‘bacterial,’ ‘viral,’ or ‘noninfected’ were generated for all 193 patients. In the main chart review, physicians were presented with host response results for only the 100 patients in the ‘test cohort’. The host response calibration chart review questions, and additional details about methods are available in Supplementary Text.

### Chart Review

We recruited three physicians with subspecialty training in infectious diseases to perform a retrospective chart review on the 200 patients in a blinded manner. They were asked to classify the infection status of the patients and the clinical relevance of mNGS and VirCapSeq-VERT results, in a staged fashion: 1) with only medical charts; 2) with the addition of mNGS and VirCapSeq-VERT results; and 3) with the further addition of host response results. Details are provided in Supplementary Text--Appendix S1, and Appendix S2.

Data from mNGS, VirCapSeq-VERT, host response, and physician chart reviews for all 200 patients are provided in Supplementary Data 1.

## RESULTS

### Patient population

We recruited 200 consecutive patients in the ED with suspected sepsis; applied mNGS, VirCapSeq-VERT, and host response profiling on blood specimens of each patient; and evaluated patient clinical records in two separate physician chart reviews (Fig. 1).

**Figure 1.**
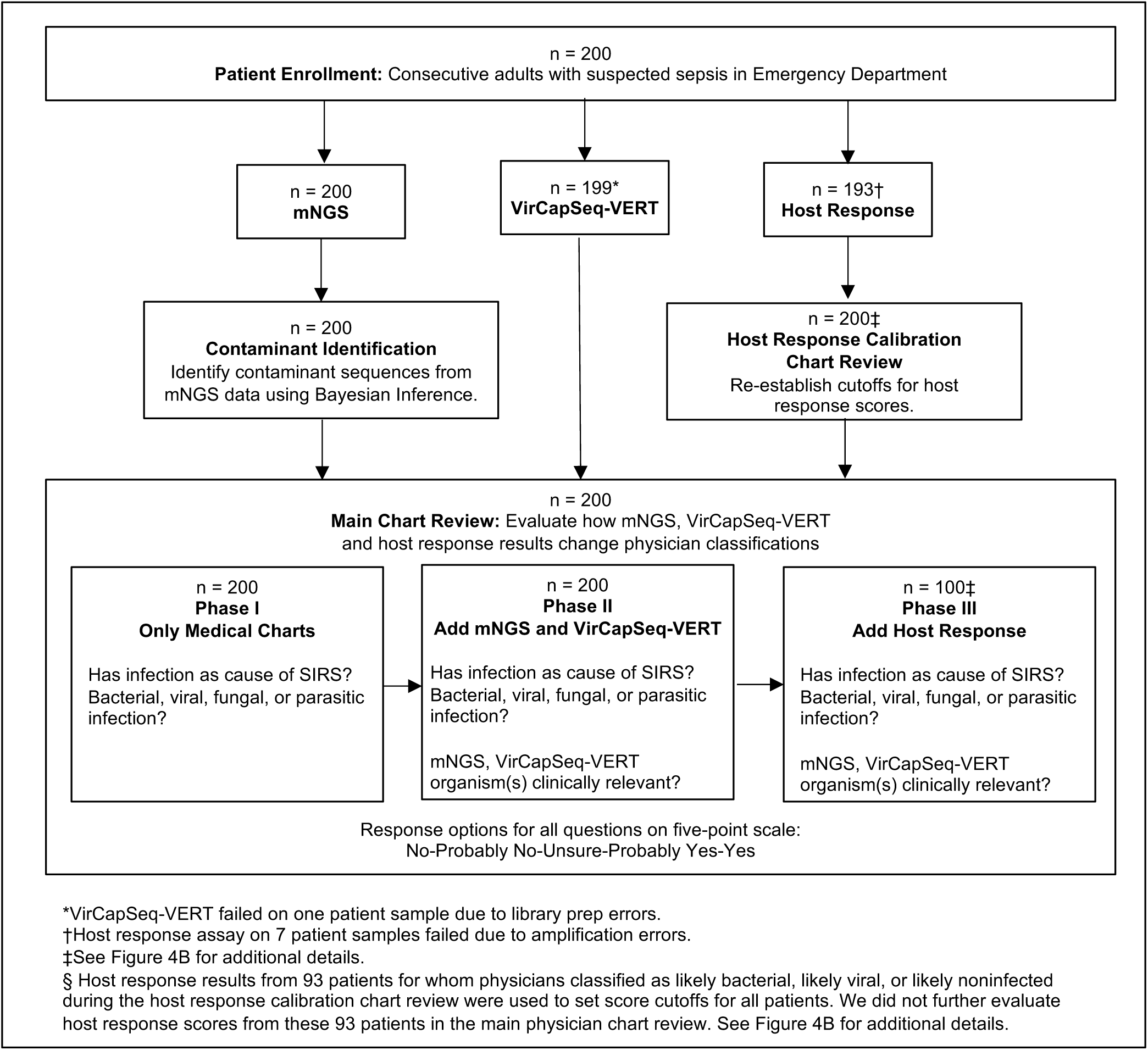
Study design. We applied three diagnostic approaches to a cohort of 200 consecutive adult patients with suspected sepsis: 1) direct bacterial DNA detection and characterization in plasma with mNGS, and contaminant sequence identification using Bayesian inference; 2) direct viral DNA and RNA enrichment and detection with viral capture sequencing (VirCapSeq-VERT) in plasma; and 3) transcript-based host response profiling with a previously-defined 18-gene assay in whole blood. Additionally, two separate chart reviews were performed. First, a ‘Host Response Calibration Chart Review’ established baseline diagnoses of all patients for the purpose of calibrating host response cutoffs. Second, a ‘Main Chart Review’ evaluated changes in diagnostic decision-making among three expert physicians by unblinding the three molecular test results in a staged fashion.

The clinical syndromes at presentation were diverse and included fever without localizing findings (32% of patients), as well as syndromes involving the respiratory (21.5%) and genitourinary (9.5%) tracts, and intra-abdominal sites (16.5%) (Table S1). Even though these patients met SIRS criteria and were suspected of having sepsis at the time of presentation, physicians classified 16 of them (8%) as not infected during the main chart review while blinded to mNGS, VirCapSeq-VERT, and host response profiling results. The remaining patients were classified as having bacterial (69 patients, 34.5%), viral (11 patients, 5.5%), fungal or co-infections (4 patients, 1%), or probable infection or unsure status (100 patients, 50%) (Fig. 2). Changes in physician classifications after considering mNGS, VirCapSeq-VERT, and host response results are summarized in Fig. 2.

**Figure 2.**
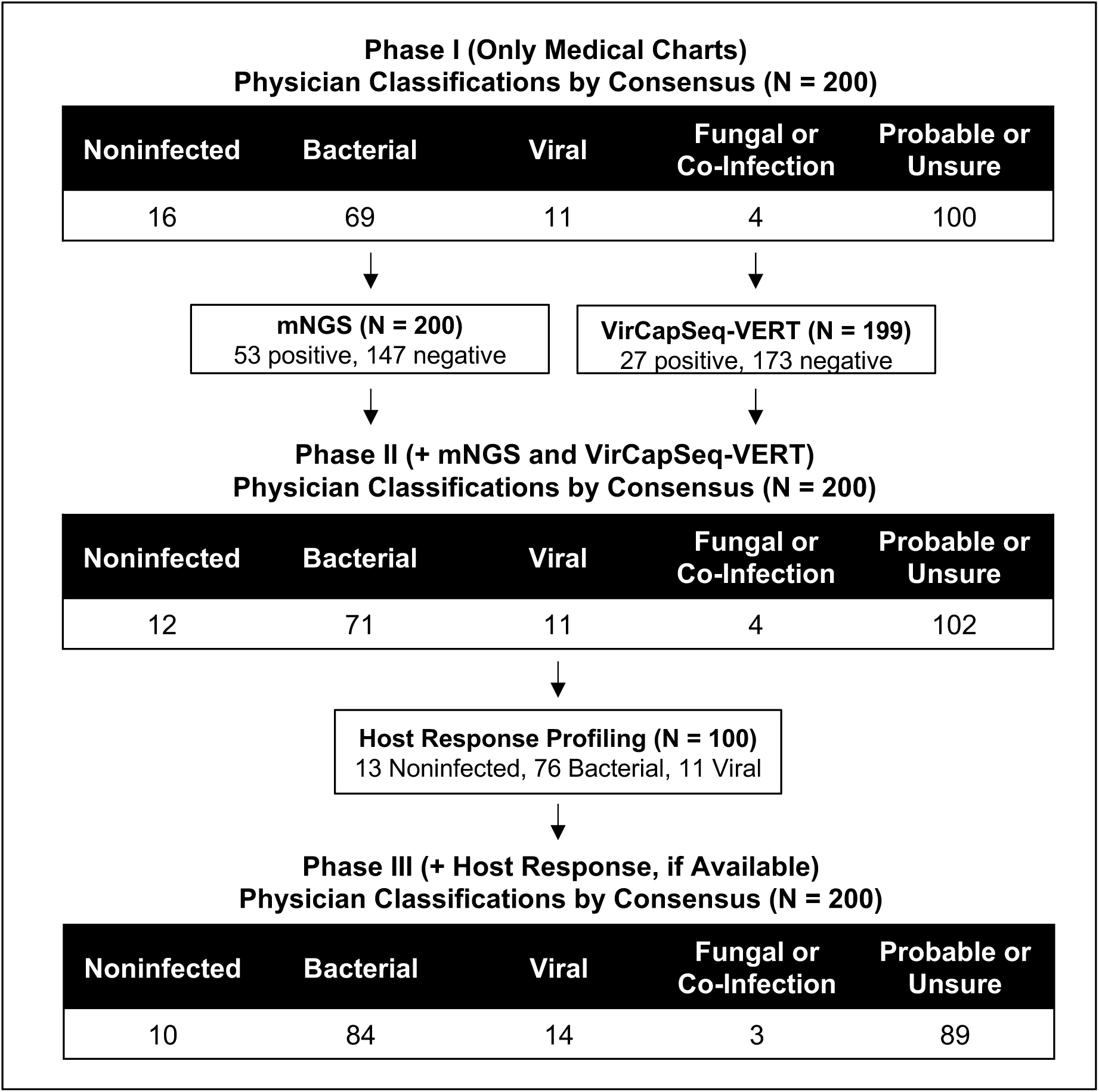
Introduction of mNGS, VirCapSeq-VERT, and host response profiling led to changes in physician classifications. At each phase of the main chart review, patients were assigned with high confidence to one of four diagnostic categories or classified with only a probable diagnosis (e.g., probable bacterial, probable noninfected) or unsure diagnosis by a panel of three physicians. Physicians did not evaluate host response scores from seven patients in whom the assay failed due to amplification errors, and from 93 patients who had host response scores used to set cutoffs. The same classification from Phase II was used for patients in Phase III who did not have host response scores for evaluation.

### Comparison of mNGS and VirCapSeq-VERT with standard-of-care microbiology

To distinguish signal from noise and remove contaminant sequences from plasma sequence data, we developed a gamma-Poisson mixture model-based Bayesian inference method. Using the 40 negative control samples, this method identified the vast majority of taxa in our dataset as contaminants (Fig. S3). Subsequent analyses of mNGS output were performed on contaminant-filtered data.

Bacterial sequences were identified by mNGS in plasma matching those of the species cultivated from blood collected at the same time, from the same subject in 14 of 26 patients with positive blood cultures (Table S2). Interestingly, mNGS results were also concordant with the positive results of urine or sputum cultures performed within 1 day of presentation in 3 of 24 patients with negative blood cultures. To test whether sequencing depth might explain low sensitivity, we selected plasma samples from 7 patients with positive blood, urine, wound, or bronchoalveolar lavage cultures but negative mNGS results and acquired an additional 65-262 million reads per sample. With these additional data, 2-111 sequencing reads matching the species of the isolated organism(s) were recovered in 6 of the 7 samples (Table S3).

VirCapSeq-VERT high-throughput sequencing was performed on 199 of the 200 available plasma samples, generating an average of 12 million reads per sample. One of the 200 samples failed to yield sufficient nucleic acid for analysis despite repeated extraction attempts. VirCapSeq-VERT analysis of plasma detected cytomegalovirus DNA in 2 of 2 subjects, but did not reveal viruses that were subsequently identified by PCR with respiratory and stool samples from 8 patients, as well as by a heterophile antibody (Monospot) Epstein-Barr Virus test in 1 patient performed within 1 day of presentation (Table S2). Respiratory or stool samples were not tested using VirCapSeq-VERT and there were no independent molecular or culture data indicative of viremia.

Of the 40 patients with organisms detected by mNGS in plasma from the day of presentation that were not identified with standard-of-care microbiological testing performed within 5 days of presentation, organisms in 14 patients were classified by physician consensus as either ‘probably clinically relevant’ or ‘clinically relevant’ (Fig. 3, Table 1). The addition of mNGS results led physicians to change their classification for the presence of bacterial infection in just six of these 14 patients by consensus (Fig. 3). The remaining eight patients were already established as known bacterial infection patients by physician consensus before mNGS results were revealed.

**Table 1.**
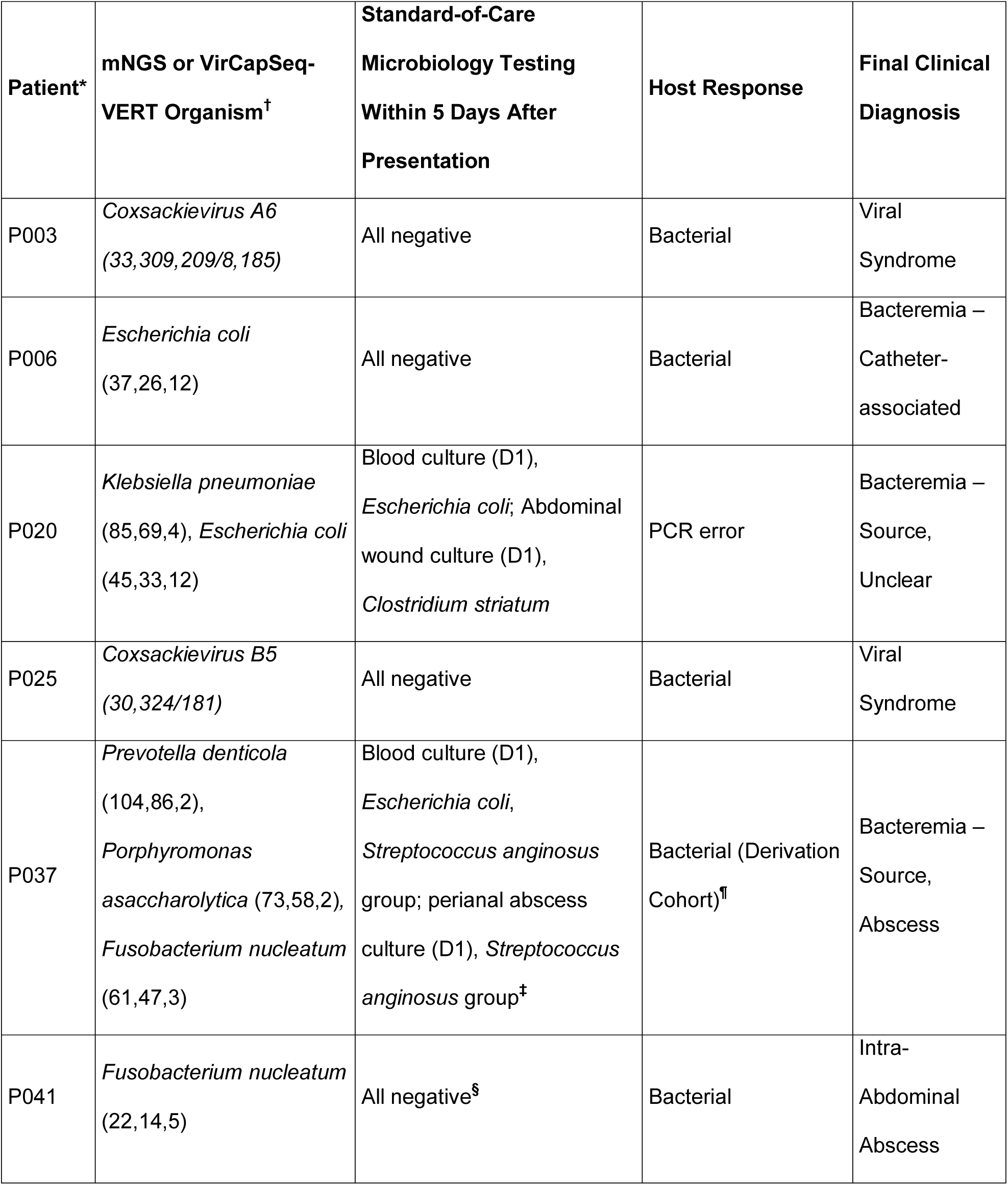
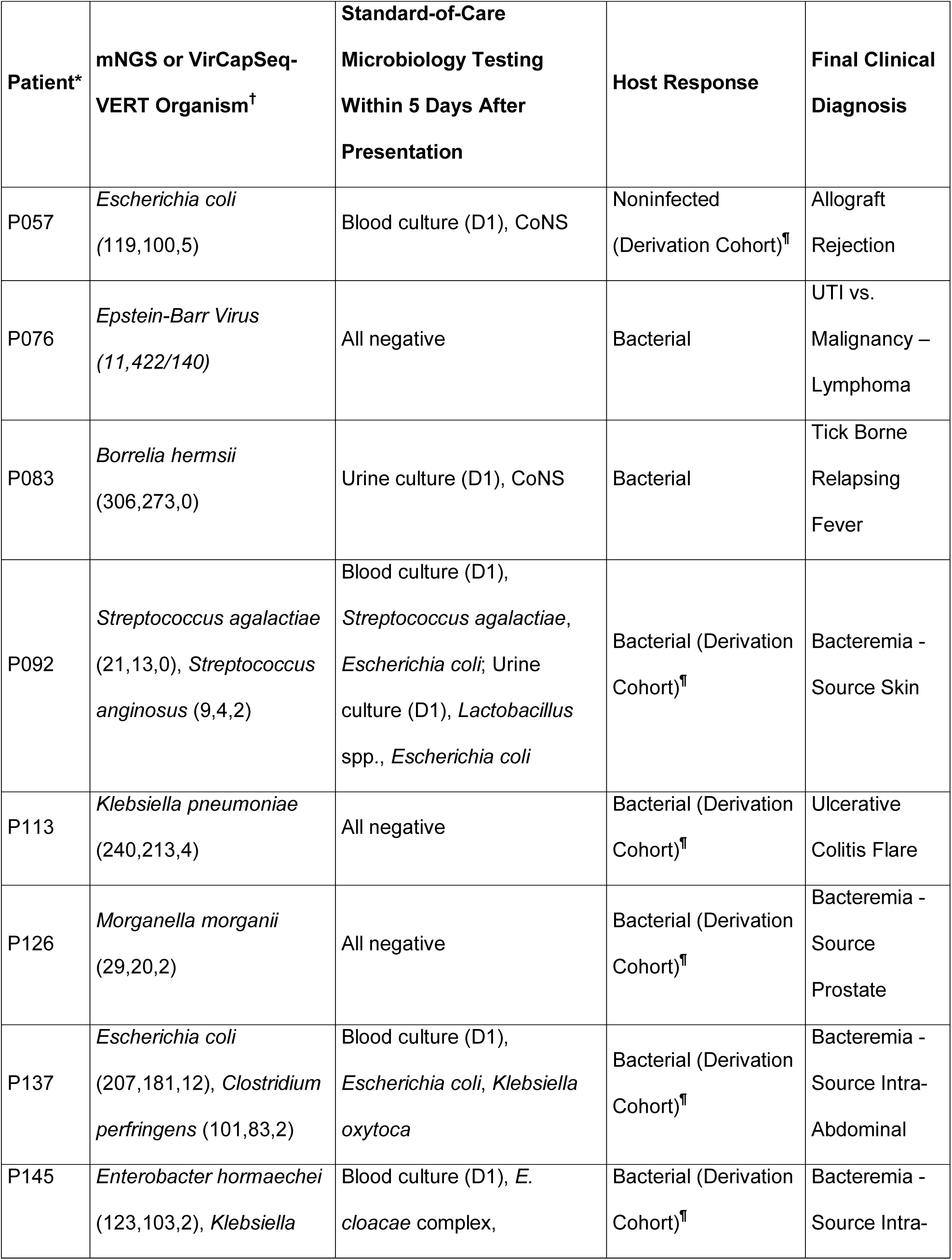
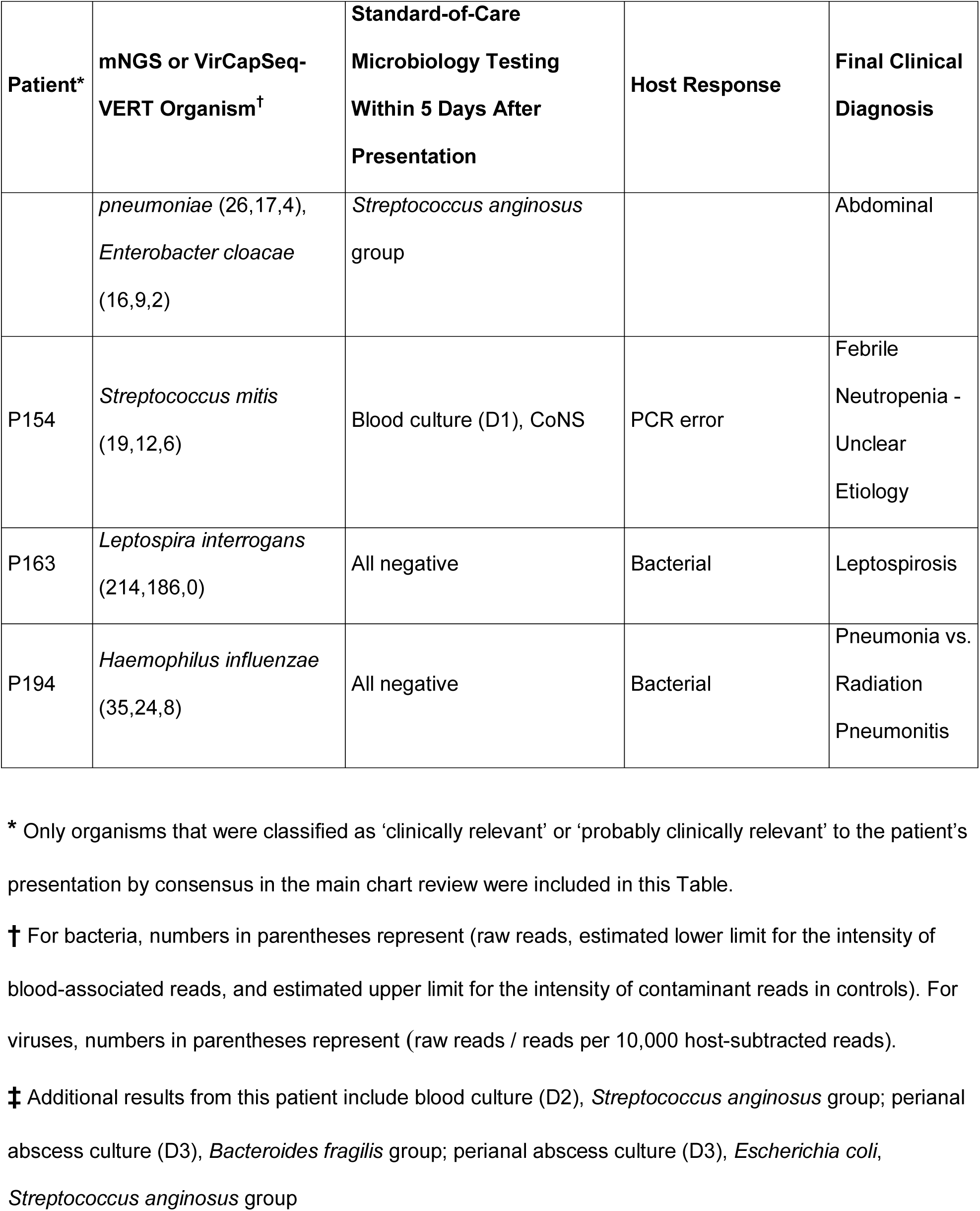
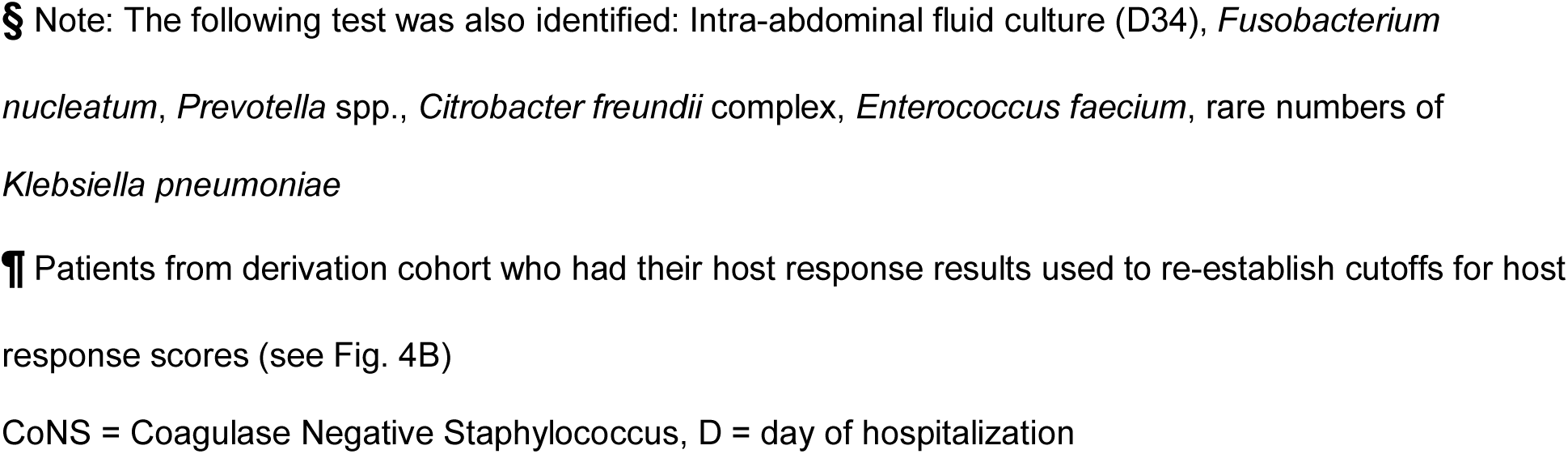
Patients with ‘clinically relevant’ or ‘probably clinically relevant’ organism(s) detected by mNGS or VirCapSeq-VERT, and not detected by hospital tests

| **Patient\*** | **mNGS or VirCapSeq-**<br>**VERT Organism<sup>†</sup>** | **Standard-of-Care**<br>**Microbiology Testing**<br>**Within 5 Days After**<br>**Presentation** | **Host Response** | **Final Clinical**<br>**Diagnosis** |
| --- | --- | --- | --- | --- |
| P003 | *Coxsackievirus A6*<br>(33,309,209/8, 185) | All negative | Bacterial | Viral<br>Syndrome |
| P006 | *Escherichia coli*<br>(37,26,12) | All negative | Bacterial | Bacteremia –<br>Catheter-<br>associated |
| P020 | *Klebsiella pneumoniae*<br>(85,69,4), *Escherichia coli*<br>(45,33,12) | Blood culture (D1),<br>*Escherichia coli*; Abdominal<br>wound culture (D1),<br>*Clostridium striatum* | PCR error | Bacteremia –<br>Source,<br>Unclear |
| P025 | *Coxsackievirus B5*<br>(30,324/181) | All negative | Bacterial | Viral<br>Syndrome |
| P037 | *Prevotella denticola*<br>(104,86,2),<br>*Porphyromonas*<br>*asaccharolytica* (73,58,2),<br>*Fusobacterium nucleatum*<br>(61,47,3) | Blood culture (D1),<br>*Escherichia coli*,<br>*Streptococcus anginosus*<br>group; perianal abscess<br>culture (D1), *Streptococcus*<br>*anginosus* group<sup>‡</sup> | Bacterial (Derivation<br>Cohort)<sup>¶</sup> | Bacteremia –<br>Source,<br>Abscess |
| P041 | *Fusobacterium nucleatum*<br>(22,14,5) | All negative<sup>§</sup> | Bacterial | Intra-Abdominal<br>Abscess |

| Patient* | mNGS or VirCapSeq-VERT Organism<sup>†</sup> | Standard-of-Care<br>Microbiology Testing<br>Within 5 Days After<br>Presentation | Host Response | Final Clinical<br>Diagnosis |
| --- | --- | --- | --- | --- |
| P057 | *Escherichia coli*<br>(119,100,5) | Blood culture (D1), CoNS | Noninfected<br>(Derivation Cohort)<sup>¶</sup> | Allograft<br>Rejection |
| P076 | *Epstein-Barr Virus*<br>(11,422/140) | All negative | Bacterial | UTI vs.<br>Malignancy –<br>Lymphoma |
| P083 | *Borrelia hermsii*<br>(306,273,0) | Urine culture (D1), CoNS | Bacterial | Tick Borne<br>Relapsing<br>Fever |
| P092 | *Streptococcus agalactiae*<br>(21,13,0), *Streptococcus*<br>*anginosus* (9,4,2) | Blood culture (D1),<br>*Streptococcus agalactiae*,<br>*Escherichia coli*; Urine<br>culture (D1), *Lactobacillus*<br>spp., *Escherichia coli* | Bacterial (Derivation<br>Cohort)<sup>¶</sup> | Bacteremia -<br>Source Skin |
| P113 | *Klebsiella pneumoniae*<br>(240,213,4) | All negative | Bacterial (Derivation<br>Cohort)<sup>¶</sup> | Ulcerative<br>Colitis Flare |
| P126 | *Morganella morganii*<br>(29,20,2) | All negative | Bacterial (Derivation<br>Cohort)<sup>¶</sup> | Bacteremia -<br>Source<br>Prostate |
| P137 | *Escherichia coli*<br>(207,181,12), *Clostridium*<br>*perfringens* (101,83,2) | Blood culture (D1),<br>*Escherichia coli*, *Klebsiella*<br>*oxytoca* | Bacterial (Derivation<br>Cohort)<sup>¶</sup> | Bacteremia -<br>Source Intra-Abdominal |
| P145 | *Enterobacter hormaechei*<br>(123,103,2), *Klebsiella* | Blood culture (D1), *E.*<br>*cloacae* complex, | Bacterial (Derivation<br>Cohort)<sup>¶</sup> | Bacteremia -<br>Source Intra- |
|  | *pneumoniae* (26,17,4),<br>*Enterobacter cloacae*<br>(16,9,2) | *Streptococcus anginosus*<br>group |  | Abdominal |
| P154 | *Streptococcus mitis*<br>(19,12,6) | Blood culture (D1), CoNS | PCR error | Febrile<br>Neutropenia -<br>Unclear<br>Etiology |
| P163 | *Leptospira interrogans*<br>(214,186,0) | All negative | Bacterial | Leptospirosis |
| P194 | *Haemophilus influenzae*<br>(35,24,8) | All negative | Bacterial | Pneumonia vs.<br>Radiation<br>Pneumonitis |

\* Only organisms that were classified as 'clinically relevant' or 'probably clinically relevant' to the patient's presentation by consensus in the main chart review were included in this Table.
† For bacteria, numbers in parentheses represent (raw reads, estimated lower limit for the intensity of blood-associated reads, and estimated upper limit for the intensity of contaminant reads in controls). For viruses, numbers in parentheses represent (raw reads / reads per 10,000 host-subtracted reads).
‡ Additional results from this patient include blood culture (D2), *Streptococcus anginosus* group; perianal abscess culture (D3), *Bacteroides fragilis* group; perianal abscess culture (D3), *Escherichia coli*, *Streptococcus anginosus* group
§ Note: The following test was also identified: Intra-abdominal fluid culture (D34), *Fusobacterium nucleatum*, *Prevotella* spp., *Citrobacter freundii* complex, *Enterococcus faecium*, rare numbers of *Klebsiella pneumoniae*
¶ Patients from derivation cohort who had their host response results used to re-establish cutoffs for host response scores (see Fig. 4B)
CoNS = Coagulase Negative Staphylococcus, D = day of hospitalization

**Figure 3.**
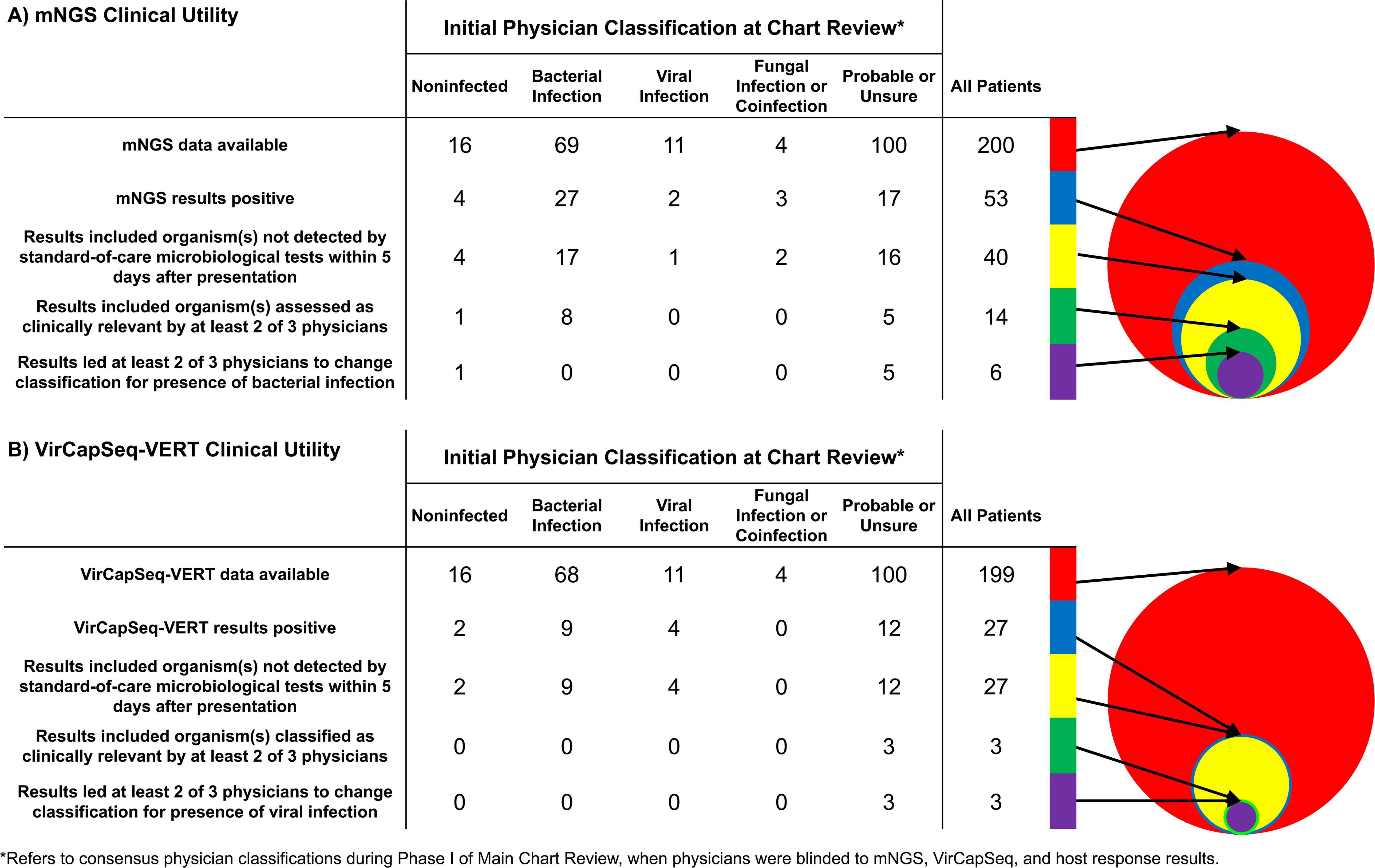
Clinical utility of positive mNGS and VirCapSeq-VERT results. Tables and Venn diagrams illustrate the number of patients with clinically relevant mNGS and VirCapSeq-VERT results which reveal the etiologies of patient presentations and change diagnostic decision-making. Patients were grouped according to positive (A) mNGS and (B) VirCapSeq-VERT results for microorganisms that 1) were not detected by standard-of-care microbiology performed within 5 days after presentation, 2) were classified as ‘clinically relevant’ or probably ‘clinically relevant’ to the patient’s presentation by physicians blinded to host response results during the main chart review, and 3) led physicians to change their classification of the patient for bacterial/viral infection by at least 1 point on a 5-point Likert scale. Patients were assigned to one of five groups using consensus classifications made during the main chart review when physicians were blinded to mNGS, VirCapSeq, and host response results. Details for all patients with positive mNGS and VirCapSeq-VERT results for microorganisms or viruses that were not detected by standard-of-care microbiological testing are presented in Tables S4 and S6, respectively.

**Figure 4.**
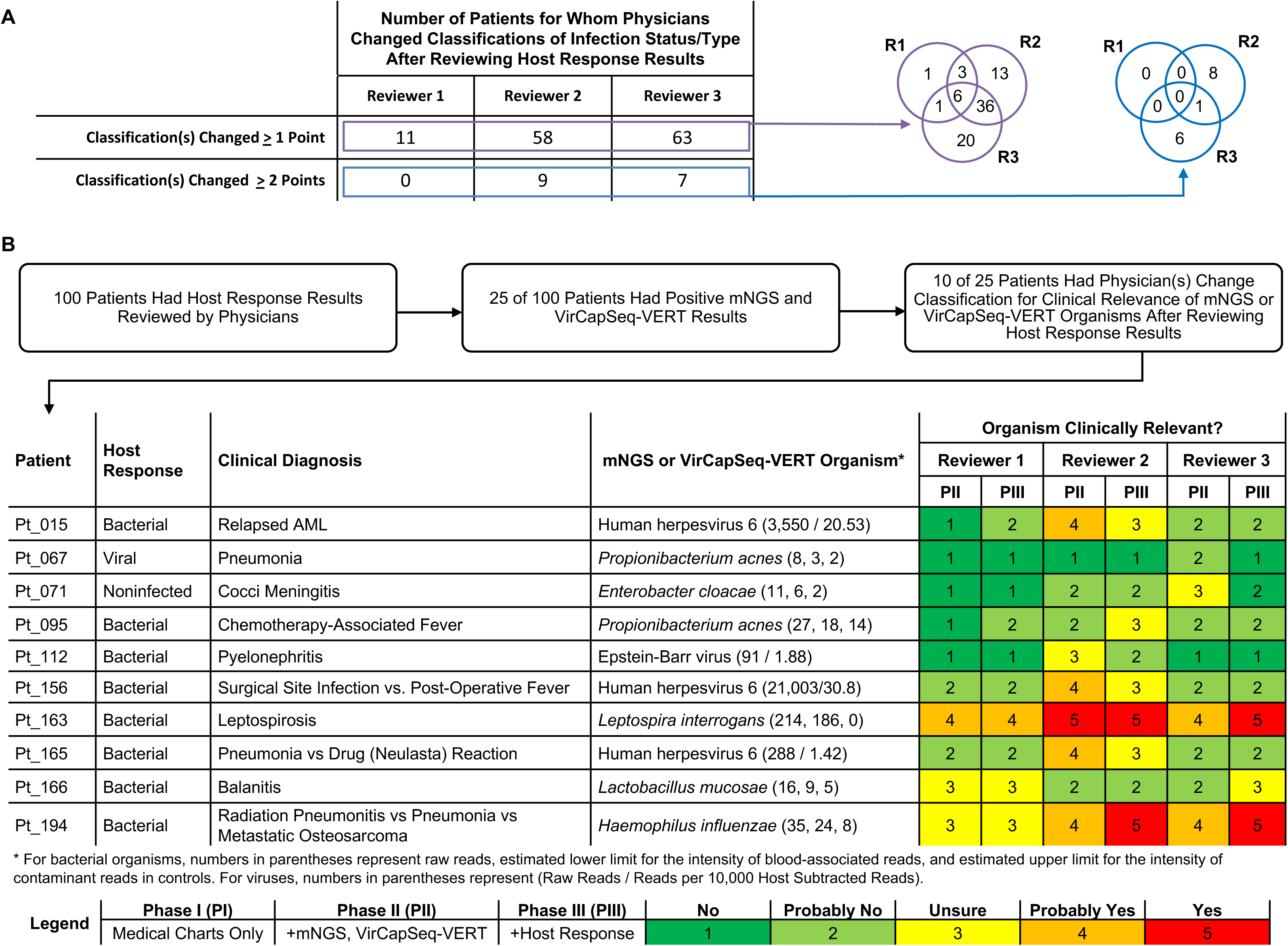
Clinical utility of host response results. In the main chart review, physicians considered the host response profiling results of the 100 patients in the validation cohort. (A) Table and Venn diagrams illustrate the number of patients for whom the introduction of host response results led physicians to change their classification(s) of infection status and type by at least one or two points. Response options were on a five-point scale (No-Probably No-Unsure-Probably Yes-Yes). (B) Twenty-five of the 100 test cohort patients had positive mNGS or VirCapSeq-VERT results. After reviewing host response profiling results, physicians changed their assessment of the clinical relevance of the putative infectious agent to the patient’s presentation in 10 of the 25 patients. The assessments for these 10 patients before (Phase II, PII) and after (Phase III, PIII) unblinding of the physicians to the host response profiling results are presented in this table.

The putative bacterial agents found only by mNGS included *Streptococcus mitis* (P154), *Borrelia hermsii* (P083), *Leptospira interrogans* (P163), and *Haemophilus influenzae* (P194) (Table 1). In five patients with positive blood cultures (P020, P037, P092, P137, and P145), mNGS revealed organisms that were not found in the blood cultures (Table 1). mNGS results and clinical details for all 40 patients with organisms not detected by standard-of-care microbiological testing are presented in Table S4.

Of the 27 patients with viruses detected by VirCapSeq-VERT that were not identified with standard-of-care microbiological tests, viruses in three of them were classified by physician consensus as either ‘probably clinically relevant’ or ‘clinically relevant’ to the clinical presentation, and led to a change in patient classification by the physician reviewers (Fig. 3, Table S5, with clinical details for each patient presented in Table S6). Two of these patients had Coxsackievirus sequences, and one had probable Epstein-Barr Virus infection. VirCapSeq-VERT demonstrated utility in detecting potential chronic viral infection or viral reactivation in 18 patients with human herpesvirus 6, hepatitis C virus, hepatitis B virus, BK virus, Epstein-Barr virus, or Trichodysplasia spinulosa-associated polyomavirus. The remaining virus sequences had uncertain clinical relevance (Tables S5, S6).

Among the patients for whom physician classifications of infection status and type were most altered by the results of mNGS and VirCapSeq-VERT (Table S7), P163 was a male traveler to South Asia who returned with headache, fever and diarrhea, and was initially assessed by treating physicians to have a viral infection but had sequences in plasma that matched *Leptospira interrogans*. P083 presented with fever after spending time in the Sierra mountains, was presumed to have a urinary tract infection by treating physicians, but was found with mNGS to have tick-borne relapsing fever due to *Borrelia hermsii*.

### Impact of Host Response Profiling Results on Physician Classifications

To evaluate the impact of host mRNA response signatures on physician classifications of patients, we applied the previously-established Integrated Antibiotics Decision Module (IADM)^25^ on 193 patient samples. The IADM incorporates the Sepsis MetaScore (SMS) which distinguishes noninfection- and infection-associated SIRS, and the Bacterial/Viral metaScore (BVS) which distinguishes bacterial and viral infections (Fig. S4A). Using cutoffs established with a ‘derivation cohort’ of 93 patients adjudicated by physicians in a separate chart review (Fig. S4B), host response classifications of ‘bacterial,’ ‘viral,’ or ‘noninfected’ for all 193 patients were compared to physician assessments (Fig. S4C, S4D). We then examined the impact of host response profiling results on physician diagnostic decision-making of the 100 test cohort patients who were not used in setting host response score cutoffs. In 46 patients (46%), the addition of host response profiling results led at least two of three physicians to change their classification of infection status and type (Fig. 5A). We also asked physicians to assess the clinical relevance of organisms detected with mNGS and VirCapSeq-VERT using medical charts only, and then with the addition of host response results. Ten patients had at least one physician change their assessment of clinical relevance of an organism revealed by mNGS or VirCapSeq-VERT upon receiving host response scores (Fig. 5B).

### Possible bacterial DNA bloodstream translocation in patients originally classified as noninfected

In plasma samples from eight of the 50 patients originally assessed by physician consensus as probably noninfected or noninfected, mNGS detected sequences of bacteria associated with the human indigenous microbiota (Table S8). Physicians noted pre-existing mucosal membrane disturbances in five of these eight patients, thus raising the possibility of bacterial DNA translocation from heavily colonized mucosal sites. For example, P070, who had high abundances of sequences from more than 20 oral cavity-associated organisms in plasma, also had documented gingivitis and hemoptysis. All eight patients improved after their ED visit, six of whom were not prescribed antibiotics. Host response results could have been useful to physicians for interpreting ambiguous mNGS results from these patients. However, most of these patients were in the ‘derivation cohort’ used for setting host response cutoffs, and thus did not have their host response results assessed (Fig. S4B). Nonetheless, host response profiling predicted that five of the eight patients were not infected (Table S8).

## DISCUSSION

Diagnosing infections in patients with suspected sepsis is challenging, particularly in those with multiple co-morbidities. We applied two broad-range sequencing approaches, mNGS and VirCapSeq-VERT, as well as host response profiling to a prospectively-sampled cohort of 200 adults with suspected sepsis, representative of real-world patient heterogeneity in a tertiary care hospital ED. We evaluated diagnostic decision-making by three infectious disease physicians as they received information from the electronic medical record, the two sequencing-based methods, and host response profiling in a staged fashion. Our results showed that sequencing methods can detect clinically relevant organisms that are missed by routine microbiological diagnostic methods, as well as other organisms that may not be clinically relevant.

Our mNGS positivity rate was comparable to those cited in other clinical metagenomics studies^35,36^. For example, in a study of 204 meningitis and encephalitis patients, diagnoses in 13 of them were made solely by metagenomic sequencing with CSF samples, with an impact on patient management in 7 of the 13^7^. In a study of cell-free plasma in 358 febrile sepsis patients, 15% of patients had probable causal pathogens detected solely by metagenomic sequencing^5^. In our study, results for 9 of the 17 patients with detected organisms that were otherwise missed by routine testing led reviewing physicians to change their classifications of infection status and type. Contaminant sequence identification and computational removal represents one of the greatest barriers to expanding the clinical application of metagenomic sequencing, especially in specimens with low microbial biomass such as blood. The gamma-Poisson mixture model-based Bayesian inference approach that we have introduced here offers an important advance over other published methods in addressing this challenge.^4,37,38,39^

As metagenomic sequencing enters clinical practice, it is important to recognize the potential of this powerful approach to reveal both clinically-relevant and -irrelevant microbial sequences. There are multiple reasons for, and sources of the latter, including translocation of microbial nucleic acids from heavily colonized body sites, reactivation of latent viruses, and contamination of laboratory reagents or specimen collection devices. Clinicians are accustomed to the importance of clinical-pathological correlations for establishing the relevance of laboratory findings. But with the advent of sensitive molecular diagnostic technologies, this challenge will only grow. In addressing this challenge, we illustrate the utility of transcriptional host response signatures, as an objective adjunct in guiding the interpretation of mNGS results and avoiding misdiagnosis and unnecessary treatment. Our results add to those of Langelier et al., who combined host response and metagenomic sequencing to diagnose lower respiratory tract infections^31^ using a different approach from ours. In addition to the use of signatures trained and validated on thousands of patients^25^, we demonstrated the clinical utility of data integration with expert physician case reviews.

The major limitation of our mNGS protocol was suboptimal sensitivity, which was explained in part by the choice of sequencing depth. Low numbers of patients with known systemic viral infections limited our ability to assess the performance characteristics of VirCapSeq-VERT. Bioinformatic errors^17^ from contaminant, misannotated, or missing genomes in microbial databases, and other technical limitations such as index-hopping^40^, may have led to false-negative and false-positive findings. Recent efforts to curate databases^41^, manufacture contaminant-free extraction kits, and enrich for microbial sequences^6,8^ are steps in the right direction to prepare metagenomic sequencing for routine clinical use, but much work remains to be done.

The host response profiling assay classified many viral and noninfected patients as bacterial. One possible reason for these misclassifications was the strict dichotomous cutoffs that we used to distinguish infected versus noninfected cases, and viral versus bacterial infections. Reporting results with numeric values rather than dichotomous cutoffs will allow better weighting of these scores in patient assessments. In addition, further work is needed to establish and lock cutoffs, validate on additional patient populations, and quantify test characteristics.

The measurement of a diagnostic tool’s ability to change clinical decision-making, rather than just a comparison of its results to standard-of-care testing, is a valuable component of establishing clinical utility. Our proof-of-concept study on a consecutive, prospectively-sampled patient cohort suggests that integrating host response profiling with metagenomic sequencing may synergistically enhance the utility of each assay, and ultimately, the diagnosis of patients with suspected sepsis.

## Acknowledgements

We thank Ian Brown, Patrice Callagy, Cheryl Bucsit, and Adele Araya of the Stanford Emergency Department for facilitating the collection of samples for this project. We thank members of the Relman Lab, and in particular, Stephen J. Popper, Eitan Yaffe, Christine L. Sun, Daniela S. Goltsman, and Natalie Campen for valuable advice, feedback, and general assistance. We also thank members of the Lipkin Lab, Joel A. Garcia, Nishit P. Bhuva, and Lokendrasingh Chauhan for technical assistance, and Bohyun Lee and Komal Jain for bioinformatics support, Kelly Murphy (Stanford Emergency Department) for advice and assistance, and John Coller and Xuhuai Ji at the Stanford Functional Genomics Facility for their technical assistance with the Fluidigm platform. We thank Alvaro Hernandez and Chris Wright at the High-Throughput Sequencing and Genotyping Unit at the University of Illinois at Urbana-Champaign for their assistance and support with DNA library preparation and sequencing.

