## Supplemental Materials for "Combined use of metagenomic sequencing and host response profiling for the diagnosis of suspected sepsis"

### Supplementary Material

#### Table of Contents

**Supplementary Text** **3**

**Methods** **3**

**Discussion** **24**

**Appendix S1.** Host Response, mNGS, and VirCapSeq-VERT interpretation information provided to physician chart reviewers **29**

**Appendix S2.** Questions in main physician chart review **36**

**Figure S1.** Contribution of batch effects to sample composition profiles **44**

**Figure S2.** Two distinct reagent contamination profiles in negative controls **45**

**Figure S3.** Application of Bayesian inference for distinguishing blood-associated DNA sequences from contaminating DNA sequences in mNGS data **47**

**Figure S4.** Host response calibration **49**

**Figure S5.** Receiver operating characteristic curves for host response scores **51**

**Table S1.** Clinical characteristics of patient population **52**

**Table S2.** Performance of plasma mNGS and VirCapSeq-VERT versus clinically-indicated standard-of-care microbiology **54**

**Table S3.** Additional bacteria identified after sequencing seven plasma samples to higher depth **56**

**Table S4.** Clinical relevance of all microorganisms found by mNGS that were not detected by hospital tests **59**

**Table S5.** Clinical relevance of all viruses found by VirCapSeq-VERT that were not detected by hospital tests: Summary Table **77**

**Table S6.** Clinical relevance of all viruses found by VirCapSeq-VERT that were not detected by hospital tests: Clinical details **79**

**Table S7.** Clinical details for four cases in which mNGS and VirCapSeq-VERT had the greatest influence in changing physician classifications **89**

**Table S8.** Physician interpretation and host response results for patients originally classified as noninfected or probably noninfected, and found by mNGS to have bacterial sequences in plasma **92**

**Supplementary References** **96**

### Supplementary Methods

#### Subject Enrollment

This study was approved by the Stanford University Administrative Panel on Human Subjects Research (Protocols 32851, 29803, and 29733). The patient cohort was a prospective, consecutive convenience sample of 200 patients with suspected sepsis. Plasma and PAXgene™ RNA whole blood samples were prospectively collected from adult patients presenting to the Stanford University Hospital Emergency Department (ED) who were not pregnant; met 2 of 4 SIRS criteria, as defined by Bone et al.;[1] and were suspected to have infection at presentation by triage nurses or other clinicians.

Blood samples for this study were collected at the time of venipuncture for standard-of-care bacterial cultures during presentation under a waiver of informed consent granted by the IRB. From this patient sample bank, we then identified 200 consecutive suspected sepsis patients (spanning a 128-day period in 2016) who met the following additional criteria:

1) 2.5 mL of whole blood in a PAXgene RNA tube (collected as part of this study protocol) and at least 200 μL of plasma, were available;

2) blood samples underwent nucleic acid extractions without errors; and

3) no access restrictions for their electronic medical record.

We note that our patient sample banking operations began before the new sepsis-3 definition was released in 2016.[2] Thus, our enrollment efforts did not include patients that would have otherwise been identified under the expanded sepsis-3 definition.

In addition, we collected 2.5 ml of peripheral blood in a PAXgene RNA tube from each of 10 healthy adult volunteers in the San Francisco Bay Area to serve as controls for host response profiling. Written, informed consent was obtained from each healthy volunteer prior to sampling.

**Healthy Volunteer Subjects**

*Inclusion Criteria*

1) Must be healthy adults between the ages of 18 and 75 years, and should not be under the care of a physician for any chronic condition

2) Must be able to read, understand and sign the approved consent form

3) Must be able and willing to follow study procedures and instructions

4) Must be willing and able to transport stool samples from their home to the Stanford Center for Translational & Clinical Research

*Exclusion Criteria*

1) Have used systemic or intra-oral antibiotics or antifungals within the 6-month period preceding study enrollment

2) Require antibiotics before dental treatments

3) Have fewer than 15 teeth

4) Are pregnant

5) Have a condition known to compromise the immune system, including HIV infection

Criteria regarding stool samples, antimicrobials, and oral health were included because the healthy volunteer samples in this study were also used for a concurrent study of bacterial translocation of microbial sequences in blood during states of health.

Of our 10 healthy volunteers, 6 were female. The median age was 32 years, with an interquartile range of 26.75 to 35.5 years.

**mNGS Plasma Processing and Sequencing**

Whole blood samples were obtained from 200 suspected sepsis patients in 6 mL EDTA tubes by nurses and phlebotomists at the Stanford ED, and immediately stored at 4$^{o}$C. Plasma was prepared from whole blood by centrifuging (1,500 x g for 10 minutes) within 72 hours of collection and stored at -80C. Prior to extraction, plasma samples underwent an additional centrifugation step at 16,000 x g. DNA was extracted from 400 µL of plasma with the QIAGEN Circulating Nucleic Acid extraction kit. (In some patients, only 200-400 µL of plasma was available due to low blood draw volume.) Three negative controls (molecular-grade water drawn into an EDTA tube) were included in every extraction batch of 24 samples, for a total of 36 negative controls across all batches. Extracted DNA was quantified with the Quant-iT dsDNA Assay Kit, high sensitivity (ThermoFisher Scientific), and quality-controlled with the AATI Fragment Analyzer or 2100 Bioanalyzer using the Agilent High Sensitivity D1000 kit.

Libraries were prepared with the KAPA HyperPrep Kit (Roche) at the High-Throughput Sequencing and Genotyping Unit at the University of Illinois at Urbana-Champaign. The starting amount of plasma DNA ranged from 4.96 ng to 0.669 ng, with one outlying sample at 1.542 ng. All libraries were prepared using the Kappa Hyper Kit

(Roche) without size selection and sequenced on the HiSeq 4000 (Illumina) with 2x150 nucleotide paired-end reads.

In a pilot experiment, we sequenced DNA from a plasma sample from each of 15 patients with a positive blood culture, at a depth of 40-60 million reads/sample, and from 4 negative controls at a depth of 2-6 million reads/sample, using unique single-indexed 10-nt barcodes. Because of the potential impact of barcode hopping[3] on sequencing data, we used unique dual-indexed barcodes for library prep for the remaining plasma samples from the other 185 patients, and obtained 10-52 million reads/sample, as well as 3-6 million reads/sample for each of 36 negative controls.

*Bioinformatics*. Reads were de-multiplexed with Illumina software, adapters were removed with `SeqPrep` (**https://github.com/jstjohn/SeqPrep**), low-quality bases were removed with `Sickle` (**https://github.com/najoshi/sickle**), and human reads were subtracted with `bowtie2`[4] under default parameters. `Kraken`[5] was run on all non-human reads using a database of all complete bacterial genomes and viral genomes from RefSeq, and all human, protozoa, archaeal, and fungal genomes (including contigs, scaffolds, and/or chromosomes) from RefSeq downloaded in July 2016. All genomes used in our `Kraken` database had low-complexity regions masked with DUST[6]. The human genome was added to reduce false-positive eukaryotic pathogen classifications, and a conservative filtering threshold of 0.3 was applied to reduce false-positive classifications during `Kraken` alignments. Finally, bacterial reads that were classified to the species-level by `Kraken` were imported into R and further analyzed with `phyloseq`[7].

We did not further analyze `Kraken` results for eukaryotic organisms because we found a high number of eukaryote reads across all plasma samples. We believe that the vast majority of these reads do not represent true infections but are most likely human reads that were not detected by `bowtie2`, or misalignment errors due to `Kraken` or to the RefSeq database. Additionally, we decided not to analyze further the archaea reads because of the paucity of evidence for an archaeal species acting as a human pathogen.[8]

In an exploratory analysis to identify potential batch effects, we considered a count table of 1,644 species in 200 plasma samples and 42 negative controls. Then, we reduced the number to 1,248 species by removing species that were not detected in any of the plasma samples. Next, we ranked species abundance in each sample, where the species with the largest abundance was assigned the largest rank. To reduce the artificially large difference in ranks, species with rank below a threshold of 900 were set at one.[9] PCA performed on a truncated ranking transformation showed possible batch-effects which may have contributed to variation in sample sequence composition (Fig. S1). Two distinct clusters were detected when samples were grouped into sets based on their extraction batches: Set 1, consisting of 90 plasma samples and 18 negative controls from extraction batches 1, 2, 3, 4, 11, and 12; and Set 2, which consisted of 95 plasma samples and 18 negative controls from extraction batches 5, 6, 7, 8, 9, and 10. The Pilot Set, consisting of 15 plasma samples and 6 negative controls of the pilot experiment, clustered with Set 1.

Rather than use a log scale for visualization (because of its crushing effect on intermediate-abundance species), we used `arcsinh`. We accounted for the unequal library depths using the median-of-ratios method.[10] Closer examination of species in the negative controls of Sets 1 and 2 revealed distinct contamination signatures, with numerous high-abundance taxa unique to each Set (Fig. S2). We hypothesized that differences in manufacturing lots of the nucleic acid extraction kits may have caused the variation in sample sequence composition, as Glassing et al. have previously reported.[11]

To distinguish blood-associated DNA sequences from contaminant sequences in plasma samples, we developed a Bayesian statistical method that leverages data from negative control samples.

**Computational Method for Contaminant Identification and Removal**

McMurdie and Holmes[12] proposed the use of simple gamma-Poisson mixtures (negative binomial) to model microbiome count data. Following their approach, we modelled the data generating process (factoring out the library depth effect) of species-specific reads in a plasma sample as the sum of two independent Poisson distributions that included 1) **true** reads belonging to the plasma sample, and 2) reads originating from contamination sources. Each of the Poisson distribution intensity parameters (intensity of true reads and contaminant reads) was considered to come from a gamma distribution. If we only made technological replicates with fixed library depths, we would observe a number of reads $K_{ij}$ which is the sum of two independent Poisson random variables; one with $\lambda_{ij}^{(r)}$ as the true intensity parameter and $\lambda_{i}^{(c)}$ as the contaminant intensity parameter for each species $i$ in plasma sample $j$. In reality, we observed random reads with biological variation and unequal library depth $S_j$ for each plasma sample $j$. We estimated the effect of library depth using the negative controls and median-of-ratios method that benefits from the scaling property of the gamma distribution. Based on this mixture model and our observed data, we provided a Bayesian method[13,14] for inferring the true intensity of the plasma sample microbial DNA in the presence of microbial DNA contamination using the negative controls. First, we defined a prior density for the contaminant intensity in a plasma sample for each of the species using the negative controls. Next, we found the marginal likelihood and the marginal reference prior for the true intensity in the plasma sample for each of the species. Then, using Bayesian reference analysis, we obtained the marginal posterior for the true intensities up to a constant. Finally, we used the Metropolis-Hasting (MH) Markov Chain Monte Carlo (MCMC) method to sample from the marginal posterior of the true intensity.

Then we used the Bayesian method to estimate the marginal posterior for the true intensity for a given plasma sample.

*Count matrix* $K \in \mathbb{R}^{m \times N}$

| Species | $\text{Plasma}_1$ | $\text{Plasma}_2$ | $\cdots$ | $\text{Plasma}_{n_1}$ | $\text{Control}_{(n_1+1)}$ | $\text{Control}_{(n_1+2)}$ | $\cdots$ | $\text{Control}_N$ |
|---|---|---|---|---|---|---|---|---|
| $\text{Species}_1$ | $K_{11}$ | $K_{12}$ | $\cdots$ | $K_{1n_1}$ | $K^0_{1(n_1+1)}$ | $K^0_{1(n_1+2)}$ | $\cdots$ | $K^0_{1N}$ |
| $\text{Species}_2$ | $K_{21}$ | $K_{22}$ | $\cdots$ | $K_{2n_1}$ | $K^0_{2(n_1+1)}$ | $K^0_{2(n_1+2)}$ | $\cdots$ | $K^0_{2N}$ |
| $\vdots$ | $\vdots$ | $\vdots$ | | $\vdots$ | $\vdots$ | $\vdots$ | | $\vdots$ |
| $\text{Species}_i$ | $K_{i1}$ | $K_{i2}$ | $\cdots$ | $K_{in_1}$ | $K^0_{i(n_1+1)}$ | $K^0_{i(n_1+2)}$ | $\cdots$ | $K^0_{iN}$ |

| | | | | | | | | |
|---|---|---|---|---|---|---|---|---|
| ⋮ | ⋮ | ⋮ | | ⋮ | ⋮ | ⋮ | | ⋮ |
| $\text{Species}_m$ | $K_{m1}$ | $K_{m2}$ | ⋯ | $K_{mn_1}$ | $K^0_{m(n_1+1)}$ | $K^0_{m(n_1+2)}$ | ⋯ | $K^0_{mN}$ |

The table shows the count matrix of $n_1$ plasma samples, $n_2$ negative controls and $m$ species, where $K_{ij}$ is the number of reads of species $i$ in the $j$-th plasma sample whose true prevalence we supposed to be $\mu_{ij}$ and whose dispersion parameter is $\gamma_i$, and $K^0_{il}$ is the number of reads of species $i$ in the $l$-th negative control with prevalence $\mu^0_{il}$ and dispersion $\gamma^0_i$. In notation, $K_{ij} \sim \text{NB}(d_j\mu_{ij}, \gamma_i)$ and $K^0_{il} \sim \text{NB}(d^0_l\mu^0_{il}, \gamma^0_i)$, where $d_j$ and $d^0_l$ are the linear scaling factors for plasma sample $j$ and negative control $l$ that account for the library depths $S_j$ and $S^0_l$.

If there is no contamination, the hierarchical mixture model for a plasma sample gives the number of reads $K_{ij}$ as $\text{Poisson}(\lambda_{ij}d_j)$ and $\lambda_{ij} \sim \text{gamma}\,(\alpha_{ij}, \beta_{ij})$, where $\lambda_{ij}$ is the true intensity of species $i$ in plasma sample $j$ after factoring out the library depth effect $d_j$.

In the presence of contamination, we considered the observed reads in a plasma sample to be a mixture of the true and contaminant reads. Thus, we modelled $K_{ij}$ as the sum of two independent Poisson random variables with two different intensities and we write, $K_{ij} = K^{(r)}_{ij} + K^{(c)}_{ij}$: 1) the true intensity parameter is $\lambda^{(r)}_{ij}$ and 2) the contaminant intensity parameter is $\lambda^{(c)}_{ij}$. We assumed $\lambda^{(c)}_{ij}$ follows a gamma $\left(\alpha^{(c)}_{ij}, \beta^{(c)}_{ij}\right)$ distribution, encoding our prior degree of belief of the contaminant intensity, and we wrote the contaminant parameters specific to species $i$ in each plasma sample $j$: $\alpha_{ij}^{(c)}$ and $\beta_{ij}^{(c)}$. Then, we estimated these parameters using the negative controls as in (14).

Given the plasma sample, $K_j = [K_{1j}, K_{2j}, \cdots, K_{mj}]^T$, where $\sum_{i=1}^{m} \mathbb{E}\left[K_{ij}\right] = S_j$, $\mathbb{E}\left[S_j\right] = d_j \sum_{i=1}^{m} \mu_{ij}$, and $S_j$ is the library depth of the $j$-th plasma-sample, the model for the number of reads of each species $i$ in plasma sample $j$ is written according to the hierarchical model

$$\begin{aligned} K_{ij} \mid \left(\lambda_{ij}^{(r)} + \lambda_{ij}^{(c)}\right) d_j &\sim \text{Poisson}\left(\left(\lambda_{ij}^{(r)} + \lambda_{ij}^{(c)}\right) d_j\right), \\ \Pi\left(\lambda_{ij}^{(r)}\right) &= \frac{\left|I\left(\lambda_{ij}^{(r)}\right)\right|^{1/2}}{|I(0)|^{1/2}}, \\ \lambda_{ij}^{(c)} &\sim \text{gamma}\left(\alpha_{ij}^{(c)}, \beta_{ij}^{(c)}\right), \end{aligned} \tag{1}$$

where $\Pi\left(\lambda_{ij}^{(r)}\right)$ is a marginal reference prior for the true intensity and $I(\cdot)$ is the Fisher information obtained through their marginal probability density.

Using the negative controls in Table S1, we estimated the prior density of the contaminant intensities $\lambda_{ij}^{(c)} \sim \text{gamma}\left(\alpha_{ij}^{(c)}, \beta_{ij}^{(c)}\right)$. Then, using this prior information, we derived the marginal reference prior for true intensities.

By considering the contaminant intensity as a nuisance parameter and our knowledge that $\lambda_{ij}^{(c)} \sim \text{gamma}\left(\alpha_{ij}^{(c)}, \beta_{ij}^{(c)}\right)$, the marginal model for the true intensity is

$$p\left(k_{ij}|\lambda_{ij}^{(r)}d_j\right) = \int_0^{\infty} \text{Poisson}\left(k_{ij}|\left(\lambda_{ij}^{(r)} + \lambda_{ij}^{(c)}\right)d_j\right) \text{gamma}\left(\lambda_{ij}^{(c)}|\alpha_{ij}^{(c)}, \beta_{ij}^{(c)}\right) d\lambda_{ij}^{(c)}. \qquad (2)$$

Since we can estimate the contaminant intensities as $\lambda_{ij}^{0(c)} = \frac{\alpha_{ij}^{(c)}}{\beta_{ij}^{(c)}}$ using negative controls, (2) can be simplified using a Dirac delta function

$$p\left(k_{ij}|\lambda_{ij}^{(r)}d_j\right) = \int_0^{\infty} \text{Poisson}\left(k_{ij}|\left(\lambda_{ij}^{(r)} + \lambda_{ij}^{(c)}\right)d_j\right) \delta\left(\lambda_{ij}^{(c)} - \lambda_{ij}^{0(c)}\right) d\lambda_{ij}^{(c)}$$

$$= \text{Poisson}\left(k_{ij}|\left(\lambda_{ij}^{(r)} + \lambda_{ij}^{0(c)}\right)d_j\right). \qquad (3)$$

Then we used the marginal model in (3) to compute the Fisher information:

$$\left|I\left(\lambda_{ij}^{(r)}\right)\right| = -\mathbb{E}\left[\frac{\partial^2}{\partial\left(\lambda_{ij}^{(r)}\right)^2} \log p\left(k_{ij}|\lambda_{ij}^{(r)}d_j\right)\middle|\lambda_{ij}^{(r)}\right] = \frac{1}{\left(\lambda_{ij}^{(r)} + \lambda_{ij}^{0(c)}\right)d_j}. \qquad (4)$$

Thus, the reference prior for the true intensity was

$$\Pi\left(\lambda_{ij}^{(r)}\right) = \frac{\left|I\left(\lambda_{ij}^{(r)}\right)\right|^{\frac{1}{2}}}{|I(0)|^{\frac{1}{2}}} = \sqrt{\frac{\lambda_{ij}^{0(c)}}{\lambda_{ij}^{0(c)} + \lambda_{ij}^{(r)}}}\ , \qquad (5)$$

where $\lambda_{ij}^{0(c)} = \frac{\alpha_{ij}^{(c)}}{\beta_{ij}^{(c)}}$.

By Bayes' theorem, the joint posterior density of $\lambda_{ij}^{(r)}$ and $\lambda_{ij}^{(c)}$ is

$$p\left(\lambda_{ij}^{(r)} d_j, \lambda_{ij}^{(c)} d_j | k_{ij}\right) \propto p\left(k_{ij} | \left(\lambda_{ij}^{(r)} + \lambda_{ij}^{(c)}\right) d_j\right) p\left(\lambda_{ij}^{(r)}, \lambda_{ij}^{(c)}\right)$$
$$= p\left(k_{ij} | \left(\lambda_{ij}^{(r)} + \lambda_{ij}^{(c)}\right) d_j\right) \Pi\left(\lambda_{ij}^{(c)} | \lambda_{ij}^{(r)}\right) \Pi\left(\lambda_{ij}^{(r)}\right), \quad (6)$$

where $\Pi\left(\lambda_{ij}^{(c)} | \lambda_{ij}^{(r)}\right)$ is the conditional prior density for the contaminant intensity and $\Pi\left(\lambda_{ij}^{(r)}\right)$ is the marginal prior density for the true intensity.

With the reference prior in (5), the estimate for the contaminant intensities $\lambda_{ij}^{0(c)}$, and the assumption that $\lambda_{ij}^{(r)}$ and $\lambda_{ij}^{(c)}$ are independent, the joint posterior was

$$p\left(\lambda_{ij}^{(r)} d_j, \lambda_{ij}^{(c)} d_j | k_{ij}\right) \propto p\left(k_{ij} | \left(\lambda_{ij}^{(r)} + \lambda_{ij}^{(c)}\right) d_j\right) \delta\left(\lambda_{ij}^{(c)} - \lambda_{ij}^{0(c)}\right) \sqrt{\frac{\lambda_{ij}^{0(c)}}{\lambda_{ij}^{0(c)} + \lambda_{ij}^{(r)}}}. \quad (7)$$

Hence, the marginal posterior up to a constant for the true intensities was obtained by integrating (7) with respect to $\lambda_{ij}^{(c)}$

$$p\left(\lambda_{ij}^{(r)} d_j \middle| k_{ij}\right) \propto \text{gamma}\left(\left(\lambda_{ij}^{(r)} + \lambda_{ij}^{0(c)}\right) d_j \middle| \left(k_{ij} + .5\right), 1\right). \quad (8)$$

Finally, the marginal posterior for the true intensities is

$$p\left(\lambda_{ij}^{(r)} \middle| k_{ij}\right) \propto \text{gamma}\left(\lambda_{ij}^{(r)} + \frac{\alpha_{ij}^{(c)}}{\beta_{ij}^{(c)}} \middle| \left(k_{ij} + .5\right)/d_j, 1\right) \quad \text{when} \quad k_{ij} \neq 0,$$
$$p\left(\lambda_{ij}^{(r)} \middle| 0\right) \propto \text{gamma}\left(\lambda_{ij}^{(r)} \middle| .5/d_j, 1\right) \quad \text{when} \quad k_{ij} = 0, \quad (9)$$

where $j = 1, \ldots n_1$ *and* $i = 1, \ldots, m$.

Next, we estimated $\alpha_{ij}^{(c)}$ and $\beta_{ij}^{(c)}$ in (9) using the negative controls. Then, we could sample from the marginal posterior distributions for the true intensities as in (14) by plugging in the estimates. Now we show how we estimated $\alpha_{ij}^{(c)}$ and $\beta_{ij}^{(c)}$.

We used all negative control samples to estimate the mean prevalence $\mu_{il}^{0}$, dispersion $\gamma_i^0$ and library depth scaling factor $d_l^0$ for each species $i$ in the $l$-th negative control using the negative binomial model

$$K_{il}^{0} \quad \sim \mathrm{NB}(d_l^0 \mu_{il}^0, \gamma_i^0), \quad \text{where} \quad l = 1, \dots n_2. \tag{10}$$

The negative binomial model in (10) could be written as a gamma-Poisson mixture

$$\begin{aligned} K_{il}^0 | \lambda_{il}^{(c)} d_l^0 &\sim \mathrm{Poisson}\left(\lambda_{il}^{(c)} d_l^0\right) \\ \lambda_{il}^{(c)} &\sim \mathrm{Gamma}\ (\alpha_{il}^0,\ \beta_{il}^0), \end{aligned}$$

where and $\alpha_{il}^0$ and $\beta_{il}^0$ are shape and rate parameters, respectively, so we know that

$$\alpha_{il}^0 = \frac{1}{\gamma_i^0},$$

and

$$\beta_{il}^0 = \frac{1}{\gamma_i^0 \mu_{il}^0}.$$

That is,

$$\lambda_{il}^{(c)} \sim \mathrm{gamma}\left(\frac{1}{\gamma_i^0}, \frac{1}{\gamma_i^0 \mu_{il}^0}\right). \tag{11}$$

We removed the library depth effect using the median-of-ratios method by computing

$$d_l^0 = \text{median}_{i:\overline{K_\iota} \neq 0} \frac{K_{il}^0}{\overline{K_\iota}}, \tag{12}$$

where $\overline{K_\iota} = \left(\prod_{l=1}^{n_2} K_{il}^0\right)^{1/n_2}$. We estimated $\gamma_i^0$ using the three-step procedure in Love et. al. (2014)[15] that depends both on the library depth scaling factor $d_l^0$and mean prevalence of species $i$ in negative control $l$, $\mu_{il}^0$.

To define the prior density for the contaminant intensity in a plasma sample from $\lambda_{il}^{(c)} \sim \text{gamma}\left(\frac{1}{\gamma_i^0}, \frac{1}{\gamma_i^0 \mu_{il}^0}\right)$, we assumed that contamination is plasma sample-dependent only through the library depth scaling factor, i.e., $\frac{\lambda_{il}^{(c)}}{d_j} d_l^0 = \lambda_{ij}^{(c)}$.

Using the scaling property of the gamma distribution (that only changes the shape parameter), we obtained the prior density for the contaminant intensity in plasma sample j as

$$\lambda_{ij}^{(c)} \sim \text{gamma}\left(\frac{d_l^0}{d_j}\frac{1}{\gamma_i^0}, \frac{1}{\gamma_i^0 \mu_{il}^0}\right). \tag{13}$$

From (13) we chose $l$ that gives the median of $\frac{d_l^0 \mu_{il}^0}{d_j}$, where $l = 1, \ldots, n_2$.

Now we can write $\alpha_{ij}^{(c)}$and $\beta_{ij}^{(c)}$

$$\alpha_{ij}^{(c)} = \frac{d_l^0}{d_j}\frac{1}{\gamma_i^0} \quad \text{and} \quad \beta_{ij}^{(c)} = \frac{1}{\gamma_i^0 \mu_{il}^0}. \tag{14}$$

We plugged in $\hat{\alpha}_{ij}^{(c)}$ and $\hat{\beta}_{ij}^{(c)}$ to the formula in (9) and used MCMC to sample from the marginal posterior for the true intensity.

Finally, we could compute the 95% highest posterior density interval for the true intensity $\left(L_{ij}^{(r)}, U_{ij}^{(r)}\right)$ and 95% highest density interval for the contaminant intensity $\left(L_{ij}^{(c)}, U_{ij}^{(c)}\right)$ for each different species in a plasma sample. Species with a lower limit $L_{ij}^{(r)}$ smaller than the upper limit $U_{ij}^{(c)}$ were identified as contaminants. This meant that there was a 95% chance that the species was a contaminant.

An open-source R package of this method is available at **https://github.com/PratheepaJ/BARBI**. We ran the contaminant removal method on each batch, Set 1, Set 2, and the Pilot Set, with the observed abundance data. We analyzed the Pilot Set separately, even though it behaved similarly to Set 1, because the two sets were extracted by separate technicians and sequenced using different barcode adapters several months apart.

**VirCapSeq-VERT High Throughput Sequencing**

Plasma samples (150 μl) were mixed with NucliSens buffer and total nucleic acid extracted on the easyMag instrument (bioMerieux). Ten microliters of extract were subjected to reverse transcription with random hexamer priming (SuperScript III, Thermo Fisher) and second strand DNA synthesis with Klenow fragment polymerase (New England Biolabs). The resulting cDNA/DNA preparation was fragmented by sonication to an average size of 250 bp (E210 sonicator, Covaris), purified (AxyPrep), and up to 50 ng sheared product (Qubit) used for library preparation with KAPA kits (Hyper Library Preparation kit, KAPA Biosystems) and custom dual uniquely indexed barcode adaptors (Integrated DNA Technologies). The libraries were evaluated for quality and quantity by TapeStation (4200 System, Agilent) and pooled at equimolar quantities for hybridization with the ~2 million oligonucleotides comprising the VirCapSeq-VERT biotinylated probe library (47°C, O/N; NimbleGen/Roche). Each pool included a negative control (Salmon nucleic acid) that had been processed alongside the samples in the pool. Sequences hybridized to biotinylated probes were collected by magnetic streptavidin beads (DynaMag-2 magnet; Thermo Fisher), washed, and on-bead amplified by low cycle post-hybridization PCR (SeqCap EZ accessory kit V2; NimbleGen/Roche). Amplification products were purified (Agencourt Ampure beads; Beckman Coulter) and quantitated (TapeStation) for sequencing on HiSeq 2500 sequence analyzer (Illumina).

*Bioinformatics analysis.* Sequence reads were demultiplexed with Illumina software, Q30-filtered, and further cleaned by PRINSEQ v20.2[16]. Sequence data were then depleted of host background by alignment to human reference sequences downloaded from the NCBI database. Host-depleted reads were *de novo* assembled with MIRA v4.0[17] and the resulting contigs as well as remaining unique singletons subjected to homology search by MegaBlast against the NCBI non-redundant nucleotide database. Sequences that showed poor or no homology at the nucleotide level were screened by BLASTX against the viral protein database, and subsequently the whole database to exclude forced alignments and potentially false positives. Based on BLAST results the best matching NCBI sequence entries were identified and downloaded as scaffolds for mapping the entire data set to recover partial or complete genome sequences (`Bowtie2` mapper 2.0.6; http://bowtie-bio.sourceforge.net). `SAMtools` v0.1.19[18] were used to generate consensus genomes and coverage statistics. `Geneious` (v10; www.geneious.com) or `Tablet`[19] were employed to visualize and evaluate read mappings.

Read yields were normalized to 10,000 host-depleted total reads and a positive viral signal was assigned to samples with a normalized read count >0.2 after subtraction of reads occasionally recorded in the negative Salmon nucleic acid control and for which these reads distributed to at least three genome regions.

#### Host RNA transcript profiling

We tested samples from 193 patients and 10 healthy adult volunteers with a previously described 18-gene host-response assay consisting of 1) an 11-gene set to distinguish noninfection- and infection-associated SIRS, the Sepsis MetaScore (SMS)[20]; and 2) a 7-gene set to distinguish bacterial and viral infections, the 'bacterial-viral metascore' (BVS)[21]. The Stanford Functional Genomics Facility extracted RNA from PAXgene RNA tubes using the QIAcube system (Qiagen) according to the manufacturer's recommendations, and then performed qRT-PCR for specific human transcripts in triplicate using commercial TaqMan assays on the Biomark HD platform (Fluidigm). Samples from seven patients were not profiled because of failure of PCR amplification. SMS and bacterial-viral scores were calculated as previously described[21].

Since this was the first use of qRT-PCR to measure these target mRNAs, we needed to re-establish SMS and BVS cutoffs for the data generated in this study. First, physicians with subspecialty training in infectious diseases (not the three physicians of the main chart review) conducted a ‘host response calibration chart review’ to establish baseline classifications of infection status and type for the 193 patients with host response results. Each patient’s medical records were reviewed by two physicians who were blinded to mNGS, VirCapSeq-VERT, and host response profiling results. SMS and BVS score cutoffs were then re-established using the results from the ‘derivation cohort’ of 93 patients who were adjudicated as noninfected, or as having a bacterial or viral infection by physicians with evidence from standard-of-care microbiological tests. With these score cutoffs, host response classifications of ‘bacterial,’ ‘viral,’ or ‘noninfected’ were generated for all 193 patients. Among derivation cohort patients, bacterial, viral, and noninfection detection sensitivities were 93.8%, 45.5%, and 43.8% respectively. Bacterial, viral, and noninfection detection specificities were 70.4%, 97.5%, and 94.7%, respectively. The IADM distinguished noninfected and infected patients with an area under curve (AUC) of 0.73 (95% confidence interval [CI] 0.68-0.79), and bacterial from viral infections with an AUC of 0.89 (95% CI 0.85–0.93) (Fig. S4) using receiver operating characteristic (ROC) analysis (Fig. S5).

In the main chart review, physicians were presented with plots of host response results incorporating score cutoffs for only the 100 patients in the ‘test cohort’. The host response calibration chart review results are available in Supplementary Data 1.

**Host Response Calibration Chart Review**

Four physicians with subspecialty training in infectious diseases performed retrospective physician chart reviews to establish likely admission diagnoses while being blinded to sequencing and host response assay results. Each physician was assigned to 100 patients such that two physicians reviewed each patient. If there were discrepancies among the two physicians' classifications, both physicians met in person to discuss and adjudicate the classification. The following questions were reviewed during this first chart review:

1. Infection status at time of enrollment?
    a. Answer choices: Yes, Possible, No
2. Lab evidence of a clinically significant **bacterial** infection from specimen taken within first 5 days after enrollment? (Select yes if patient w/lab evidence is possibly infected.)
    a. Answer choices: Yes, No.
3. Lab evidence of a clinically significant **viral** infection from specimen taken within first 5 days after enrollment? (Select yes if patient w/lab evidence is possibly infected.)
    a. Answer choices: Yes, No.
4. Lab evidence of a clinically significant **fungal** infection from specimen taken within first 5 days after enrollment? (Select yes if patient w/lab evidence is possibly infected.)
    a. Answer choices: Yes, No.

5. Lab evidence of a clinically significant **parasitic** infection from specimen taken within first 5 days after enrollment? (Select yes if patient w/lab evidence is possibly infected.)
    a. Answer choices: Yes, No.

All questions were in regard to the patient's presentation, i.e. SIRS or sepsis.

**Physician Chart Review**

After obtaining sequencing and host response assay results, the main retrospective chart review was performed for all 200 patients by three additional physicians with specialty training in infectious diseases. Information provided to physicians for interpreting host response and sequencing results in this study are provided in Appendix S1. The questions in the chart review are provided in Appendix S2 and summarized below.

*Phase I:* First, physicians were provided the patient's medical record (while blinded to mNGS, VirCapSeq-VERT, and host response results), and asked to assess the following:

1. Whether the patient had an infection, and if so, a bacterial infection, viral infection, fungal infection, or parasitic infection.

*Phase II:* Next, physicians were provided with mNGS and VirCapSeq-VERT results alongside the patient's medical records, and asked to assess the following:

1. Whether the patient had an infection, and if so, a bacterial infection, viral infection, fungal infection, or parasitic infection.
2. Whether each of the patient's positive mNGS and VirCapSeq-VERT organisms, if any, were clinically relevant.

*Phase III:* Finally, physicians were provided host response results alongside the patient's medical charts, mNGS, and VirCapSeq-VERT results, and asked to assess the following:

1. Whether the patient had an infection, and if so, a bacterial infection, viral infection, fungal infection, or parasitic infection.
2. Whether each of the patient's positive mNGS and VirCapSeq-VERT organisms, if any, were clinically relevant.

For all questions, the physicians were provided with the following answer choices on a 5-point scale: Yes, Probably Yes, Unsure, Probably No, and No. All questions were in regard to the patient's presentation, i.e. SIRS or sepsis. Phase III was not conducted for the 93 patients whose host response scores were used to generate score cutoffs, nor the additional 7 patients with no host response scores due to PCR errors.

After gathering the results of Phase I, we grouped patients into categories of definite infection status (table 1 and figures 2-4) using the following guidelines:

1. Noninfected: Requires a "No" for infection question by at least two of three physicians.

2. Bacterial: Requires a “Yes” for bacterial infection question by at least two of three physicians.
3. Viral: Requires a “Yes” for viral infection question by at least two of three physicians.
4. Fungal: Requires a “Yes” for fungal infection question by at least two of three physicians.
5. Bacterial-Viral Coinfection: Requires a “Yes” for both bacterial and viral infection questions by at least two of three physicians.
6. Bacterial-Fungal Coinfection: Requires a “Yes” for both bacterial and fungal infection questions by at least two of three physicians.
7. Probable or Uncertain: Requires any choice but “No” for infection status question, and any choice but “Yes” for remaining questions by at least two of three physicians.

#### Supplementary Discussion

Diagnosing infections in patients with suspected sepsis is challenging, particularly in those with multiple co-morbidities. We applied two broad-range sequencing approaches, mNGS and VirCapSeq-VERT, as well as host response profiling to a prospectively-sampled cohort of 200 adults with suspected sepsis who were enrolled in an Emergency Department. The consecutive convenience sample set reflected real-world patient heterogeneity in a tertiary care hospital. We evaluated diagnostic decision-making by three infectious disease physicians as they received information from the electronic medical record, the two sequencing-based methods, and host response profiling in a staged fashion. Our results showed that sequencing methods can detect clinically relevant organisms that are missed by routine microbiological diagnostic methods, as well as other organisms that may not be clinically relevant. In addition, we demonstrated the potential for host response profiling to influence diagnostic decision-making and help interpret metagenomic sequencing results.

One of the most important features of unbiased, ‘shotgun’ metagenomic sequencing is that it is hypothesis-free, allowing simultaneous detection of thousands of organisms, including those difficult-to-culture. Seventeen of the 200 patients had clinically relevant organisms detected by mNGS and VirCapSeq-VERT that were not detected by standard-of-care microbiology within five days after presentation. Results from nine of these 17 patients led physicians to change their classifications of infection status and type. For example, patient Pt_083 presented with fever after travel to the

Sierra mountains and was presumed to have a urinary tract infection by treating physicians. However, this patient was determined by mNGS to have tick-borne relapsing fever due to *Borrelia hermsii*. Our positivity rate was comparable to other clinical metagenomics studies[22,23]. For example, in a study of 204 meningitis and encephalitis patients diagnoses in 13 of them were made solely by metagenomic sequencing with CSF samples, with an impact on patient management in 7 of the 13[24]. In a study of cell-free plasma in 358 febrile sepsis patients, 15% of patients had probable causal pathogens detected solely by metagenomic sequencing[25]. It should be noted that metagenomic sequencing can provide a 53-hour turnaround time[25], which is shorter than standard-of-care tests in some situations.

Contaminant sequence identification and computational removal represents one of the greatest barriers to expanding the clinical application of metagenomic sequencing, especially in specimens with low microbial biomass such as blood. The gamma-Poisson mixture model-based Bayesian inference approach that we have introduced here offers an important advance in addressing this challenge. In our implementation, we assumed that DNA sequences in plasma included those of contaminants. We then inferred the true 'intensity' of DNA sequences in a plasma sample that might be attributed to 'true' blood-associated nucleic acids. This method adds to others available to researchers for contaminant removal.[11,26] For example, for studies with fewer than three negative control samples, simply subtracting species based on their presence or abundance in negative controls[27] may be most appropriate. While the decontam[28] method is not well-suited for our data because it assumes that samples have a relatively higher biomass than controls, and that each taxon is either a contaminant or 'true' but not both, it can be very helpful for 16S rRNA gene amplicon sequence data from other kinds of samples.

As metagenomic sequencing enters clinical practice, it is important to recognize the potential of this powerful approach to reveal true signals, as well as clinically-irrelevant sequences which can occur because of translocation of microbial nucleic acids from heavily colonized body sites, reactivation of latent viruses, or contamination of laboratory reagents or specimen collection devices. Virus sequences from 18 of 27 patients with positive VirCapSeq-VERT results were associated with chronic infections or viral reactivations that were not clinically relevant to the patient's presentation. Additionally, eight of 50 patients who were originally classified as noninfected or probably noninfected had bacterial organisms detected by plasma mNGS. Five of these eight patients improved without antibiotics. Clinicians are accustomed to the importance of clinical-pathological correlations for establishing the relevance of laboratory findings. With the advent of sensitive molecular diagnostic technologies, this challenge will only grow. Indeed, Blauwkamp et al. detected organisms adjudicated as 'commensal' in 36 of 358 febrile sepsis patients (10.0%) and as 'viral reactivation' in 10 of 358 patients (2.8%) with metagenomic sequencing of cell-free plasma DNA[25]. They suggested that sequence abundance and overall clinical picture should be considered while assessing clinical relevance of metagenomics results.

Our data illustrate the utility of transcriptional host response signatures, as an objective adjunct in guiding the interpretation of mNGS results and avoiding misdiagnosis and unnecessary treatment. Our results add to those of Langelier et al., who combined host response and metagenomic sequencing to diagnose lower respiratory tract infections[29] using a different approach from ours. Their study included a sophisticated machine-learning-based integration of the complementary approaches, using a training set of 20 patients to generate signatures which identified infectious etiologies vs. commensal organisms in respiratory metagenomic sequencing results. In our study, host response signatures for identifying bacterial vs. viral vs. noninfected patients were previously trained on datasets from over 2,000 patients[21] representing a wide diversity of infectious etiologies. Additionally, we focused on measuring the clinical utility of having physician chart reviewers integrate metagenomic sequencing and host response profiling results into their clinical decision-making.

Our bacterial contaminant sequence identification method did not subtract all contaminant sequences in our mNGS dataset. Increasing the number of negative control samples in every extraction batch could aid in profiling the large diversity of contaminating taxa and thus enhance contaminant sequence removal. Furthermore, our negative control samples were only suited for identifying extraction reagent contaminants. We were not positioned to account for skin-associated contaminants or spurious sample-to-sample cross-contaminants.

The host response profiling assay classified many viral and noninfected patients as bacterial. One possible reason for these misclassifications was the strict dichotomous cutoffs that we used to distinguish infected vs. noninfected cases, and viral vs. bacterial infections. Reporting results with numeric values rather than dichotomous cutoffs will allow better weighting of these scores in patient assessments. Another reason for the misclassifications was the need to re-establish host response score cutoffs for this study's qRT-PCR platform and the small number of known viral patients with which to do so. Further work is needed to establish and lock cutoffs, validate on additional patient populations, and quantify test characteristics.

We believe that host response profiling and shotgun sequencing will soon achieve turnaround times of less than 90 minutes and 24 hours, respectively. In our chart review, physicians had access to the patients' full medical chart histories, including test results that only became available several days after presentation. If it were possible to limit the review to the first 90 minutes of each case history, we expect that there would have been greater changes in diagnostic decision-making across the stages of our chart review.

A central limitation in evaluating new diagnostic tools is the lack of a gold standard. We did our best to address this using expert physicians in a staged chart review. Fundamentally, it is impossible to determine whether the changes in patient classification were correct. However, the measurement of a diagnostic tool's ability to change clinical decision-making, rather than just a comparison of its results to standard-of-care testing, is a valuable component of establishing clinical utility. An important secondary finding was that clinicians had varying levels of trust in these new diagnostic tools. In conclusion, our proof-of-concept study on a consecutive, prospectively-sampled patient cohort suggests that integrating host response profiling with metagenomic sequencing may synergistically enhance the utility of each assay, and ultimately, the diagnosis of patients with suspected sepsis.

**Appendix S1.** Host Response, mNGS, and VirCapSeq-VERT Interpretation Information Provided to Physician Chart Reviewers

***Host Response Interpretation Information***

**Background:** The Integrated Antibiotic Decision Maker (IADM) is an 18-gene qRT-PCR host response assay that was developed by Sweeney et al.[21]. The assay consists of the 11-gene Sepsis MetaScore, which distinguishes between infection and non-infectious causes of inflammation, as well as the 7 gene Bacterial/Viral metaScore, which distinguishes between bacterial and viral infections.

**Sample Type Profiled:** Whole blood.

**Interpretation:** An example of a host response score in a patient is shown below. Two host-response scores are shown: the X axis measures the likelihood that an infection is present (as opposed to non-infectious cause of inflammation) using the SMS; the higher the score, the more likely an infection is present. The Y axis measures whether the infection is more likely bacterial or viral using the bacterial/viral metascore; the lower the score, the more likely the infection is bacterial. Note that ‘borderline’ bacterial-viral cases may indicate a weak signal, or may indicate a co-infection.

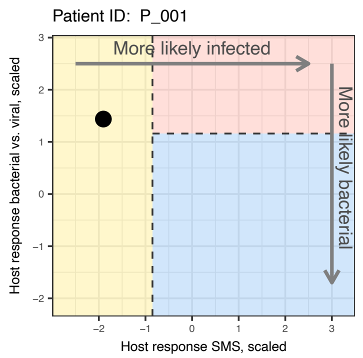

Using data from the host response calibration chart review, the cutoffs for this study were set locally to achieve a 95% sensitivity for bacterial infection when considering all three classes (bacterial, viral, non-infected).

A prior manuscript (Sweeney et al., 2016, *Science Translational Medicine*) showed that in a pooled analysis of publicly-available microarray data consisting of 1,057 samples from 20 cohorts, the IADM had 94.0% sensitivity and 59.8% specificity for bacterial infections; 53.0% sensitivity and 90.6% specificity for viral infections; and 43.0% sensitivity and 97.3% specificity for noninfectious causes of inflammation compared to retrospective chart review adjudication. The manuscript also validated the IADM on 96 pediatric patient samples using the nanoString qRT-PCR platform and showed 89.7% sensitivity and 70.0% specificity for bacterial infections; 54.5% sensitivity and 96.5% specificity for viral infections; and 61.1% sensitivity and 91.7% specificity for noninfectious SIRS.

This assay was developed from analysis of transcriptomic data using 922 adult and pediatric patients from 14 cohorts, and has so far been retrospectively validated in

2,452 patients from 38 independent cohorts. It is not yet known how this host response assay performs in complex patient populations, including immunocompromised patients.

***mNGS Interpretation Information***

**Background:** We performed metagenomic Next-Generation Sequencing (mNGS) of cell-free DNA from human plasma. DNA was sequenced to a depth of 10-70 million 2x150 nucleotide read-pairs per sample. We processed and sequenced molecular grade water in parallel as negative controls in order to characterize contaminating DNA from the lab environment and reagents. We developed and applied a statistical model to estimate and remove sequencing reads based on negative control data. Only bacterial species are presented in the final results.

**Sample Type Profiled:** DNA extracted from plasma.

**Interpretation:** The algorithm relies on the assumption that the number of raw reads of a particular species of a particular sample is equal to the sum of 1) real read-pairs that are truly present in the plasma sample, and 2) contaminating read-pairs introduced. For each species in each sample, the algorithm estimates a distribution for the number of real read-pairs, as well as a distribution for the number of contaminating read-pairs. An example of these two distributions is presented below for *E. coli* in a patient with a positive *E. coli* blood culture. Vertical red and blue bars indicating 95$^{th}$ percentile chance limits of the two estimated distributions, and the purple vertical line indicating the number of raw read-pairs. If the lower-limit of the estimated real read-pairs exceeds the upper-limit of the estimated contaminant read-pairs, the species was considered "positive" in the sample.

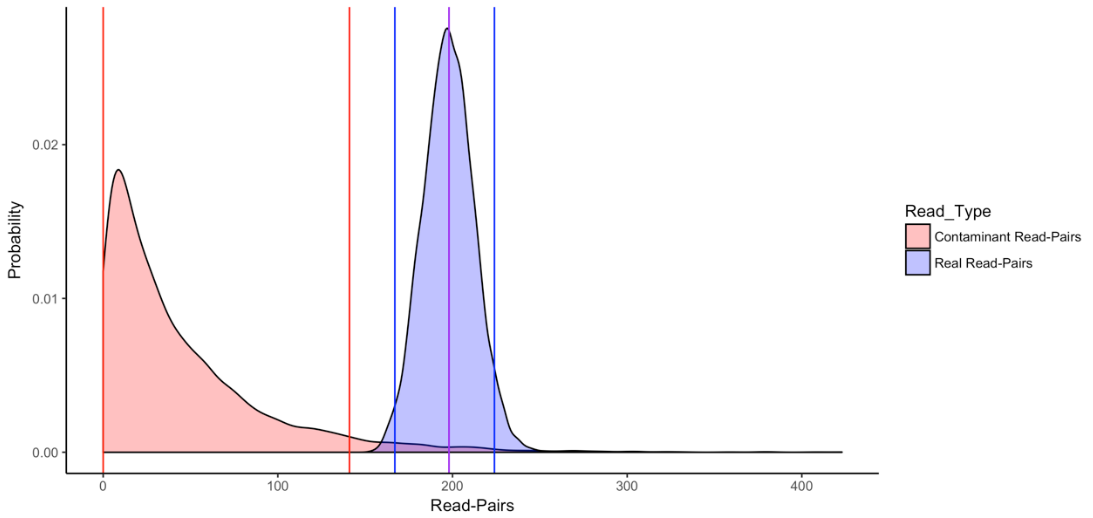

You will be provided with a table of only the positive species in each sample, along with a number of values.

An example of the positive results table from the patient mentioned above:

| | Species | Raw Reads | Lower Limit Real | Upper Limit Contaminant | In Neg. Cont. |
|---|---|---|---|---|---|
| 1 | s__Escherichia_coli | 198 | 171 | 141 | Yes |
| 2 | s__Pantoea_sp_PSNIH1 | 31 | 21 | 4 | Yes |

1. Raw reads, or the total number of reads of the species that was present in the sample.

2. Lower Limit Real, or the lower limit of the estimated real reads.

3. Upper Limit Contaminant, or the upper-limit of the estimated contaminant reads.

4. In Neg. Cont., or whether the taxon was present at all in at least one of the negative control samples.

While we made concerted efforts to control for microbial contamination, some positive organisms in the results may still represent only contaminants, as we may not have enough appropriate negative controls to gain the statistical power for perfect discrimination. Additionally, misclassifications are known to occur in NGS due to limitations in bioinformatic tools, particularly at low read counts. Thus, we suggest that you use caution in interpreting taxa if either 1) the number of raw reads, or 2) the difference between the lower-limit of the estimated real reads ("L. Real") and the upper-limit of the estimated contaminant reads ("U. Contam.") columns, are less than 5-10 reads.

Finally, because there was heavy human genome contamination and no enrichment for microbial sequences, sensitivity will be low. We recommend against relying on the negative predictive value of NGS.

*Note: The above "mNGS Interpretation Information" section was what was provided to physicians to interpret mNGS results. While writing our manuscript, we made small terminology changes:*

1. *"Estimated real/contaminant reads" are instead referred to as "true/contaminant intensity" in the rest of the manuscript.*
2. *"Distribution for the number of real read-pairs" is instead referred to as "posterior distribution for the true intensities" in the rest of the manuscript.*
3. *Read-pairs are instead referred to as "reads" in the rest of the manuscript.*

##### *VirCapSeq-VERT Information*

**Background:** VirCapSeq-VERT, or viral capture sequencing of vertebrate viruses, is a highly sensitive viral sequencing assay first introduced in 2015 by the Lipkin Lab at Columbia University (Briese et al., 2015, *mBio*). Oligonucleotide probes were used to enrich for DNA and RNA of full genomes from 207 viruses known to infect vertebrates, including humans, and enriched nucleic acids were sequenced to roughly 10 million single-ended 100 bp reads per sample.

**Sample Type Profiled:** DNA and RNA extracted from plasma.

**Interpretation:** VirCapSeq-VERT was shown in a previous study to have a 1,000 to 10,000-fold enrichment over conventional viral sequencing techniques, based on experiments with human lung tissues spiked with three respiratory viruses, and blood spiked with five viruses (Briese et al., 2015, *mBio*). Additionally, when tested on blood samples spiked with enterovirus D68, VirCapSeq-VERT showed sensitivity comparable to agent-specific PCR. Contamination is not a large problem with VirCapSeq-VERT, as the vast majority of microbial contamination is bacterial. Because of the enrichment for viral reads, we expect fewer false-positive classifications due to bioinformatic alignment errors, although they still may be possible.

Two values will be presented alongside the positive viral taxa: the raw reads, and the number of normalized reads per 10,000 host-subtracted reads. We will not present data on viruses of the family *Anelloviridae*, GB virus C, and GB virus B, all of which are not known to be pathogenic in humans.

**Appendix S2.** Questions in Main Physician Chart Review

*Note: The following is an example of a chart review for a hypothetical patient.*

#### *Introduction*

**Patient ID:** Pt_999

**MRN:** 999999999

**Date and Time of Enrollment:** 9/9/2016 09:09

**Birth Year:** 1991

**Positive hospital microbial test results within the first 5 days after enrollment:**

Urine Culture (D1): >100,000 CFU/mL E. coli

#### *Chart Review Phase I:*

**Do NOT open up the NGS, VirCapSeq-VERT, and host response results at this time.**

**What is the infection status at the time of enrollment?**

**O** Yes **O** Probably Yes **O** Unsure **O** Probably No **O** No

**Is there a clinically significant bacterial/viral/fungal/parasitic infection at the time of enrollment that is the etiology of the patient's presentation?**

| | Yes | Probably Yes | Unsure | Probably No | No |
|---|---|---|---|---|---|
| Bacterial | **O** | **O** | **O** | **O** | **O** |

| | | | | | |
|---|---|---|---|---|---|
| Viral | O | O | O | O | O |
| Fungal | O | O | O | O | O |
| Parasitic | O | O | O | O | O |

**Chart Review Phase II**

Please open up the NGS and VirCapSeq-VERT results. Do NOT open up the host response results at this time.

**For each positive VirCapSeq-VERT organism, is the organism the etiology of the patient's presentation?**

If the organism has already been identified by a hospital test in the first 5 days after enrollment, select "Already Identified by Hospital Tests."

Classify each organism in the order presented in VirCapSeq-VERT results. Leave all unnecessary fields blank. For example, if there are only two organisms, classify just VirCapSeq-VERT Organism #1 and VirCapSeq-VERT Organism #2, and leave all other fields below blank.

As a reminder, the patient's positive hospital test results (within the first five days after enrollment) are:

Urine Culture (D1): >100,000 CFU/mL E. coli

| | Already Identified by Hospital Tests | Yes | Probably Yes | Unsure | Probably No | No |
|---|---|---|---|---|---|---|
| VirCapSeq-VERT Organism #1 | o | o | o | o | o | o |
| VirCapSeq-VERT Organism #2 | o | o | o | o | o | o |
| VirCapSeq-VERT Organism #3 | o | o | o | o | o | o |
| VirCapSeq-VERT Organism #4 | o | o | o | o | o | o |
| VirCapSeq-VERT Organism #5 | o | o | o | o | o | o |

**For each positive NGS organism, is the organism the etiology of the patient's presentation?**

If the organism has already been identified by a hospital test in the first 5 days after enrollment, select "Already Identified by Hospital Tests."

Classify each organism in the order presented in NGS results. Leave all unnecessary fields blank. For example, if there are only two organisms, classify just NGS Organism #1 and NGS Organism #2, and leave all other fields below blank.

As a reminder, the patient's positive hospital test results (within the first five days after enrollment) are:

Urine Culture (D1): >100,000 CFU/mL E. coli

| | Already Identified by Hospital Tests | Yes | Probably Yes | Unsure | Probably No | No |
|---|---|---|---|---|---|---|
| NGS Organism #1 | O | O | O | O | O | O |
| NGS Organism #2 | O | O | O | O | O | O |
| NGS Organism #3 | O | O | O | O | O | O |
| NGS Organism #4 | O | O | O | O | O | O |
| NGS Organism #5 | O | O | O | O | O | O |

(Table expands for up to 25 organisms, depending on how many organisms is present in patient NGS data.)

**With the addition of NGS and VirCapSeq-VERT results, what is the infection status at the time of enrollment?**

**O** Yes **O** Probably Yes **O** Unsure **O** Probably No **O** No

**With the addition of NGS and VirCapSeq-VERT results, is there a clinically significant bacterial/viral/fungal/parasitic infection at the time of enrollment that is the etiology of the patient's presentation?**

| | Yes | Probably Yes | Unsure | Probably No | No |
|---|---|---|---|---|---|
| Bacterial | **O** | **O** | **O** | **O** | **O** |
| Viral | **O** | **O** | **O** | **O** | **O** |
| Fungal | **O** | **O** | **O** | **O** | **O** |
| Parasitic | **O** | **O** | **O** | **O** | **O** |

**Chart Review Phase II**

Please open up host response results. (Skip this section if there are no host response results.)

**With the addition of host response results: For each positive VirCapSeq-VERT organism, is the organism the etiology of the patient's presentation?**

If the organism has already been identified by a hospital test in the first 5 days after enrollment, select "Already Identified by Hospital Tests."

Classify each organism in the order presented in the VirCapSeq-VERT results. Leave all unnecessary fields blank. For example, if there are only two organisms, classify just VirCapSeq-VERT Organism #1 and VirCapSeq-VERT Organism #2, and leave all other fields below blank.

| | Already Identified by Hospital Tests | Yes | Probably Yes | Unsure | Probably No | No |
|---|---|---|---|---|---|---|
| VirCapSeq-VERT Organism #1 | O | O | O | O | O | O |
| VirCapSeq-VERT Organism #2 | O | O | O | O | O | O |
| VirCapSeq-VERT Organism #3 | O | O | O | O | O | O |
| VirCapSeq-VERT Organism #4 | O | O | O | O | O | O |
| VirCapSeq-VERT Organism #5 | O | O | O | O | O | O |

**With the addition of host response results: For each positive NGS organism, is the organism the etiology of the patient's presentation?**

If the organism has already been identified by a hospital test in the first 5 days after enrollment, select "Already Identified by Hospital Tests."

Classify each organism in the order presented in the NGS results. Leave all unnecessary fields blank. For example, if there are only two organisms, classify just NGS Organism #1 and NGS Organism #2, and leave all other fields below blank.

| | Already Identified by Hospital Tests | Yes | Probably Yes | Unsure | Probably No | No |
|---|---|---|---|---|---|---|
| NGS Organism #1 | **O** | **O** | **O** | **O** | **O** | **O** |
| NGS Organism #2 | **O** | **O** | **O** | **O** | **O** | **O** |
| NGS Organism #3 | **O** | **O** | **O** | **O** | **O** | **O** |
| NGS Organism #4 | **O** | **O** | **O** | **O** | **O** | **O** |
| NGS Organism #5 | **O** | **O** | **O** | **O** | **O** | **O** |

(Table expands for up to 25 organisms, depending on how many organisms is present in patient NGS data.)

**With the addition of NGS, VirCapSeq-VERT, and host response results, what is the infection status at the time of enrollment?**

**O** Yes **O** Probably Yes **O** Unsure **O** Probably No **O** No

**With the addition of NGS, VirCapSeq-VERT, and host response results, is there a clinically significant bacterial/viral/fungal/parasitic infection at the time of enrollment that is the etiology of the patient's presentation?**

| | Yes | Probably Yes | Unsure | Probably No | No |
|---|---|---|---|---|---|
| Bacterial | **O** | **O** | **O** | **O** | **O** |
| Viral | **O** | **O** | **O** | **O** | **O** |
| Fungal | **O** | **O** | **O** | **O** | **O** |
| Parasitic | **O** | **O** | **O** | **O** | **O** |

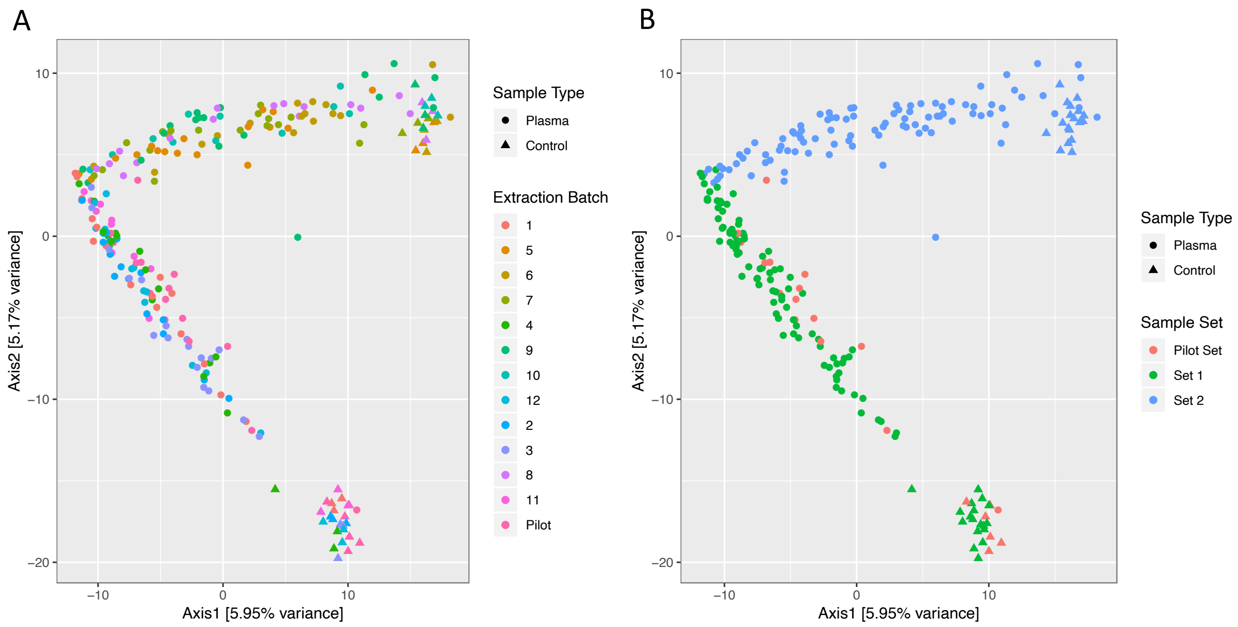

**Figure S1. Contribution of batch effects to sample composition profiles.** Principal component analysis of truncated rank-transformed plasma and negative control sample reads. (A) Individual extraction batches do not form distinct clusters. However, two distinct clusters can be visualized (B) when samples are grouped into sets based on their extraction batch. Set 1 (red) corresponds to samples in the second sequencing batch but extracted in extraction batches 1, 2, 3, 4, 11 and 12. Set 2 (green) corresponds to samples in the second sequencing batch but extracted in extraction batches 5, 6, 7, 8, 9 and 10. The Pilot Set (red) refers to all samples extracted and sequenced in the pilot sequencing batch.

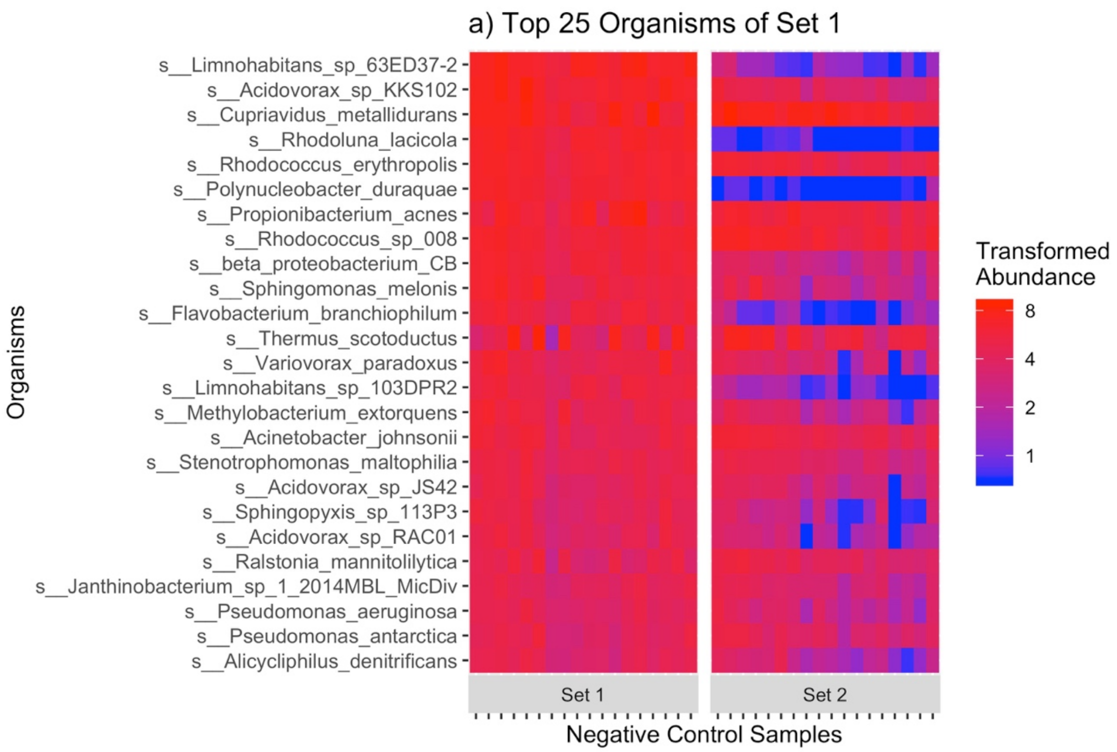
a) Top 25 Organisms of Set 1
s__Limnohabitans_sp_63ED37-2
s__Acidovorax_sp_KKS102
s__Cupriavidus_metallidurans
s__Rhodoluna_lacicola
s__Rhodococcus_erythropolis
s__Polynucleobacter_duraquae
s__Propionibacterium_acnes
s__Rhodococcus_sp_008
s__beta_proteobacterium_CB
s__Sphingomonas_melonis
s__Flavobacterium_branchiophilum
s__Thermus_scotoductus
s__Variovorax_paradoxus
s__Limnohabitans_sp_103DPR2
s__Methylobacterium_extorquens
s__Acinetobacter_johnsonii
s__Stenotrophomonas_maltophilia
s__Acidovorax_sp_JS42
s__Sphingopyxis_sp_113P3
s__Acidovorax_sp_RAC01
s__Ralstonia_mannitolilytica
s__Janthinobacterium_sp_1_2014MBL_MicDiv
s__Pseudomonas_aeruginosa
s__Pseudomonas_antarctica
s__Alicycliphilus_denitrificans
Organisms
Set 1
Set 2
Negative Control Samples
Transformed Abundance
8
4
2
1

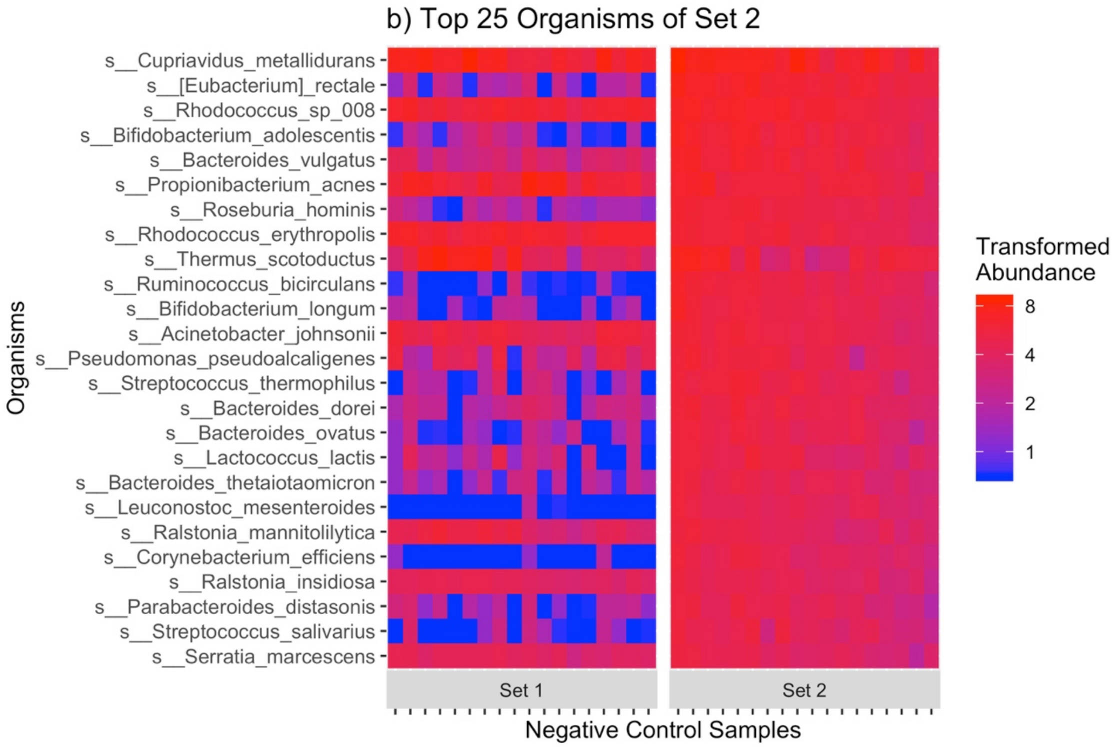
b) Top 25 Organisms of Set 2
s__Cupriavidus_metallidurans
s__[Eubacterium]_rectale
s__Rhodococcus_sp_008
s__Bifidobacterium_adolescentis
s__Bacteroides_vulgatus
s__Propionibacterium_acnes
s__Roseburia_hominis
s__Rhodococcus_erythropolis
s__Thermus_scotoductus
s__Ruminococcus_bicirculans
s__Bifidobacterium_longum
s__Acinetobacter_johnsonii
s__Pseudomonas_pseudoalcaligenes
s__Streptococcus_thermophilus
s__Bacteroides_dorei
s__Bacteroides_ovatus
s__Lactococcus_lactis
s__Bacteroides_thetaiotaomicron
s__Leuconostoc_mesenteroides
s__Ralstonia_mannitolilytica
s__Corynebacterium_efficiens
s__Ralstonia_insidiosa
s__Parabacteroides_distasonis
s__Streptococcus_salivarius
s__Serratia_marcescens
Organisms
Set 1
Set 2
Negative Control Samples
Transformed Abundance
8
4
2
1

**Figure S2. Two distinct reagent contamination profiles in negative controls.** Many of the most highly abundant taxa of Set 1 negative controls were not present in Set 2 negative controls, and vice versa. Heatmaps illustrate the arcsinh-transformed abundances of selected organisms across all Set 1 and Set 2 negative control samples prior to contaminant removal. These selected organisms consisted of a) the top 25 organisms in the negative controls of Set 1, and b) the top 25 organisms in the negative controls of Set 2, as ranked by average arcsinh-transformed abundance in negative controls of each set. A number of human commensal organisms are present as contaminants, particularly in Set 2, such as *Bifidobacterium spp.*, *Bacteroides spp.*, and *P. aeruginosa.* Set 1 and Set 2 correspond to samples in the second sequencing batch but extracted in extraction batches 1, 2, 3, 4, 11 and 12; and second sequencing batch but extracted in extraction batches 5, 6, 7, 8, 9 and 10, respectively.

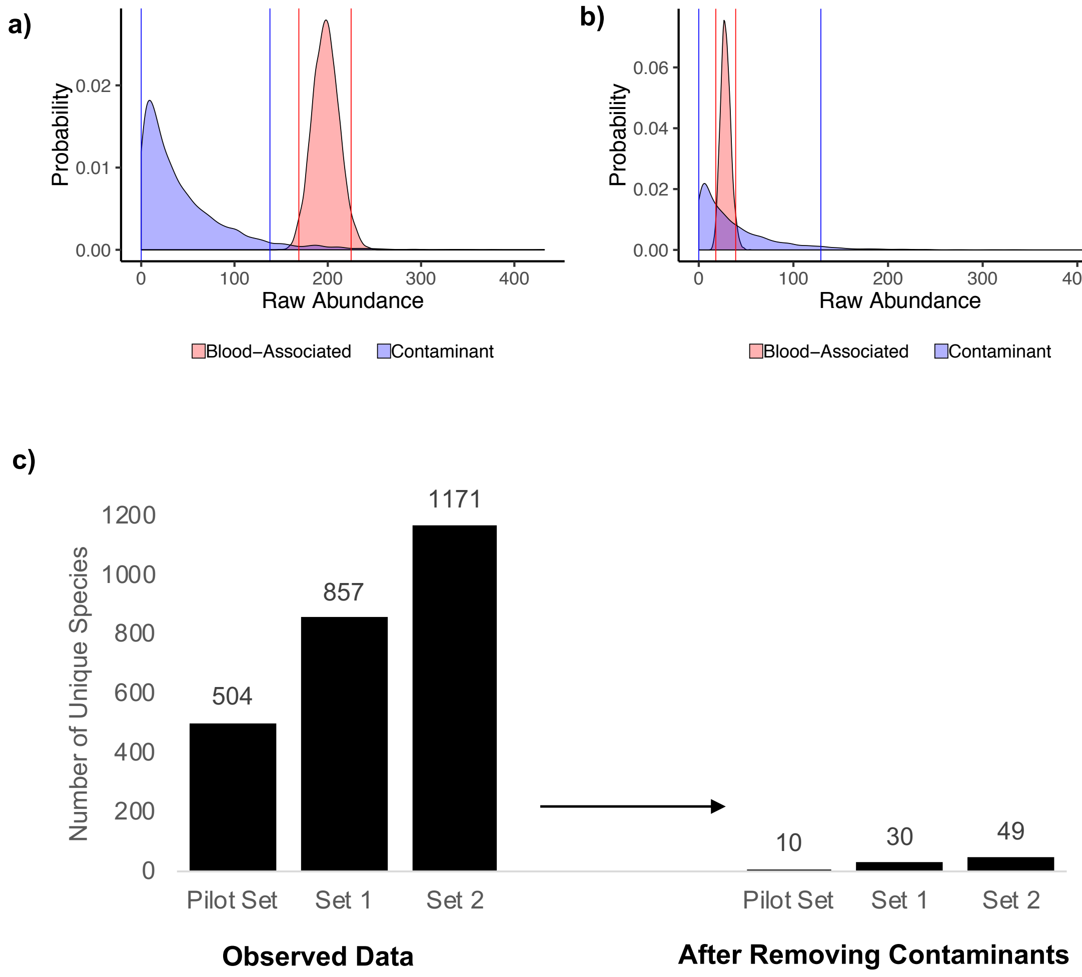

**Figure S3. Application of Bayesian inference for distinguishing blood-associated DNA sequences from contaminating DNA sequences in mNGS data.** Density plots provide the posterior distribution for true intensity and probability distribution for contaminant intensity (from negative control samples) for blood-associated *Escherichia coli* (red) and contaminating *E coli* (blue) in two patients. (A) A patient who had a positive blood culture for *E. coli* (P186), and (B) A patient who did not have a positive culture for *E. coli* (P043). Vertical lines indicate the lower and upper limits of the 95%

highest posterior density interval for true intensity (red) and highest density interval for the contaminant intensity (blue) A sample was considered to have a positive result for a particular species with 95% chance when the lower limit for the true intensity exceeded the upper limit for the contaminant intensity; otherwise, the species was eliminated from the dataset. (C) The vast majority of unique species were eliminated from samples in our three sets of plasma samples.

**A**

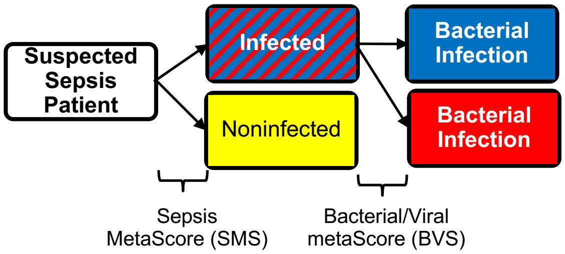

**B**

**Objective: Re-establish SMS and BVS cutoffs**

Previous cutoffs were not compatible with the qRT-PCR platform employed in this study.

n = 200

**Host Response Calibration Chart Review**

Expert physicians* established baseline diagnoses from medical charts to calibrate host response scores.

Questions: Has infection as cause of SIRS? Has bacterial, viral, fungal, or parasitic infection confirmed by hospital tests?

* Physicians different from three physicians of main chart review.

| Adjudicated Physician Classification | Number Patients* | |
|---|---|---|
| Noninfected | 32 | **Derivation Cohort (n=93)** Leverage physician classifications to train new host response score cutoffs. |
| Bacterial† | 52 | |
| Viral | 9 | |
| Bacterial-Viral Coinfection | 2 | **Test Cohort (n=100)** Present patients' host response results with new cutoffs to physicians to assess in Main Chart Review |
| Infected, Unknown Etiology | 33 | |
| Indeterminate | 65 | |

* 7 patients whose host response failed due to PCR amplification errors not included.
† Includes 3 patients with bacterial-fungal coinfections.

**C** **Example of a Patient's Host Response Result Presented to Physicians in Main Chart Review**

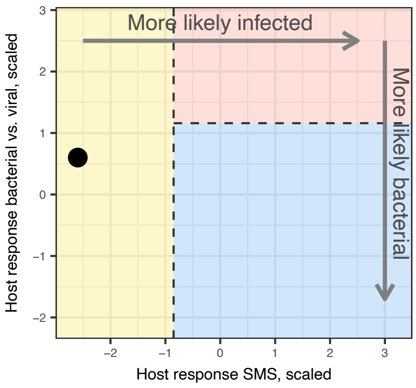

**D**

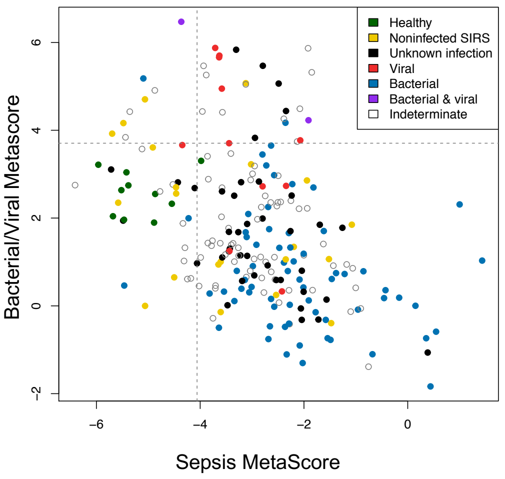

**E**

| | | | **Host Response** | | |
|---|---|---|---|---|---|
| **N = 193*** | | | **Non-Infected** | **Bact-erial** | **Viral** |
| **Adjudicated Physician Classification** | **Derivation Cohort** | **Noninfected** | 10 | 18 | 4 |
| | | **Bacterial†** | 2 | 49 | 1 |
| | | **Viral** | 1 | 4 | 4 |
| | **Test Cohort** | **Bacterial-Viral Coinfection** | 0 | 1 | 1 |
| | | **Infected, Unknown Etiology** | 3 | 29 | 1 |
| | | **Indeterminate** | 10 | 46 | 9 |

* 7 patients whose host response failed due to PCR amplification errors not included.
† Includes 3 patients with bacterial-fungal coinfections.

**Figure S4. Host response calibration.** (A) Schematic for two numeric scores of host response assay. The Sepsis MetaScore (SMS) distinguishes noninfection- and infection-associated SIRS, and the Bacterial/Viral metaScore (BVS) distinguishes bacterial and viral infections. (B) SMS and BVS score cutoffs were re-established using the results from the 'derivation cohort' of 93 of 193 patients for whom physicians adjudicated as noninfected, bacterial, or viral in a separate 'host response calibration chart review.' With these score cutoffs, host response classifications of 'bacterial,' 'viral,' or 'noninfected' were generated for all 193 patients. In the main chart review, physicians were presented with plotted host response results with score cutoffs (C) for only the 100 patients in the 'test cohort' to interpret. (C) Distribution of scores and cutoffs for the host response assay. A higher SMS indicates a higher chance of infection over noninfection, and a higher BVS indicates a higher chance of viral infection over bacterial infection. (D) Confusion matrix for the host response assay vs. adjudicated physician classifications. The following characteristics were calculated from derivation cohort patients with total n = 93: bacterial infection sensitivity, 93.9%; bacterial infection specificity, 73.2%; viral infection sensitivity, 44.4%; viral infection specificity, 93.8%; noninfected sensitivity, 31.3%; noninfected specificity, 94.8%.

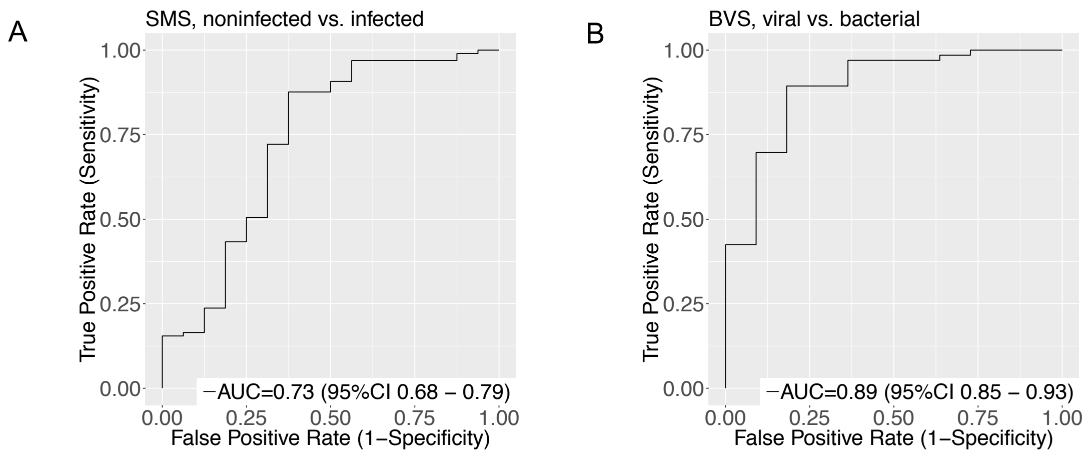

**Figure S5. Receiver operating characteristic curves for host response score.**

Receiver operating characteristic (ROC) curves for the Sepsis MetaScore (SMS, for noninfected SIRS vs. sepsis) and Bacterial/Viral metaScore (BVS).

**Table S1. Clinical characteristics of patient population.**

| | **Patients (%)** |
|---|---|
| **Infection Status and Type* (n=200)** | |
| Noninfected | 16 (8%) |
| Bacterial Infection | 69 (34.5%) |
| Viral Infection | 11 (5.5%) |
| Fungal Infection | 1 (0.5%) |
| Bacterial-Viral Coinfection | 2 (1%) |
| Bacterial-Fungal Coinfection | 1 (0.5%) |
| Probable or Unsure | 100 (50%) |
| **Type of Syndrome†** | |
| Systemic‡ | 64 (32%) |
| Respiratory | 43 (21.5%) |
| Genitourinary | 39 (19.5%) |
| Intra-Abdominal | 33 (16.5%) |
| Skin | 10 (5%) |
| Ear, Nose & Throat | 5 (2.5%) |
| Bone/Joint | 3 (1.5%) |
| Central Nervous System | 3 (1.5%) |
| **Immune Status** | |
| Immunocompromised, due to Cancer/Chemotherapy | 68 (34%) |
| Immunocompromised, due to Transplant | 15 (7.5%) |
| Immunocompromised, due to Immunosuppressing Drugs | 14 (7%) |
| Immunocompromised, due to Cancer/Chemo and Transplant | 1 (0.5%) |
| Immunocompromised, due to Congenital Disorder | 1 (0.5%) |
| Immunocompetent | 101 (50.5%) |

| | |
|---|---|
| **Neutropenia** | |
| Neutropenic | 18 (9%) |
| Not Neutropenic | 182 (91%) |
| **Admission Status** | |
| Admitted to ICU | 13 (6.5%) |
| Admitted to Floor | 134 (67%) |
| Sent Home from ED | 53 (26.5%) |
| **Sex** | |
| Female | 96 (48%) |
| Male | 104 (52%) |
| **Age (years)** | |
| Median (Interquartile Range) | 51.5 (35-68) |

*Information on infection status and type was collected from the main chart review while physicians were blinded to mNGS, VirCapSeq-VERT, and host response results. The 'probable or unsure' category includes all patients without a definite known diagnosis, including those classified as probable noninfected, probable bacterial, probable viral, and unsure. Classifications had consensus agreement by at least two of three physicians. Details on how each patient was placed into each category is provided in the Supplementary Methods. All other information in this table was extracted by a single physician at the completion of our study.

†Refers to the localization of signs and symptoms of patients at presentation.

‡Systemic refers to non-localized infection, sepsis, 'viral syndrome', fever with neutropenia, post-operative fever, and/or SIRS findings on presentation not related to infection, such as those associated with malignancies (e.g., leukemia, lymphoma, and metastatic tumors), and autoimmune disorders.

**Table S2. Performance of plasma mNGS and VirCapSeq-VERT versus clinically-indicated standard-of-care microbiology**

| | | **Standard of Care Microbiology*** | | | |
|---|---|---|---|---|---|
| | | Positive Bacterial Tests | | Positive Viral Tests | |
| | | Blood Culture (n=26) | Other Body Site Culture, PCR or immunoassay (with Negative Blood Culture)† (n=24) | Plasma PCR‡ (n=2) | Respiratory or Stool PCR, Monospot Antibody Test§ (n=9) |
| **Plasma mNGS (Bacteria) or VirCapSeq-VERT (Viruses)** | Confirmed Infectious Agent(s) | 14 | 3 | 2 | 0 |
| | No Confirmed Infectious Agent | 12 | 21 | 0 | 9 |

*Includes only tests performed within 1 day of presentation and determined to be clinically relevant to the patient's presentation by physician chart review. All blood cultures were performed at the same time as the blood draw for our study.

†Tests include the following: aerobic and/or anaerobic bacterial cultures of sputum, bronchoalveolar lavage fluid, urine, wound, and intra-abdominal abscess, and perianal abscesses; stool PCR for *Salmonella enterica* and upper throat swab PCR for *Streptococcus dysgalactiae* ssp. *Equisimilis*; and upper throat swab rapid enzyme immunoassay test for group A beta hemolytic *Streptococcus*

‡Includes plasma PCRs for CMV.

§Includes upper nasopharyngeal swab PCRs for influenza, coronavirus, respiratory syncytial virus, rhinovirus; stool PCR for norovirus; and Epstein-Barr virus Heterophile Antibody (Monospot) Test

**Table S3**. Additional bacteria identified after sequencing seven plasma samples to higher depth

| | **Raw Reads in Original Library that Revealed Bacterial Species (at 0.3 Kraken Threshold/at 0 Kraken Threshold)*** | **Raw Reads in Re-sequenced Library that Revealed Bacterial Species (at 0.3 Kraken Threshold/at 0 Kraken Threshold)*** | **Sequencing Depth in Original Library (Reads)** | **Sequencing Depth in Re-Sequenced Library (Reads)** |
|---|---|---|---|---|
| **P091 (Blood culture grew *Salmonella enterica*)** | | | 17,495,799 | 90,179,007 |
| *Salmonella enterica* | 0/18 | 5/129 | | |
| **P022 (Blood culture grew *Enterobacter cloacae* complex)** | | | 17,528,199 | 65,103,185 |

| | | | | |
|---|---|---|---|---|
| *Enterobacter cloacae complex spp.* | 0/0 | 3 /14 | | |
| **P047 (Blood culture grew CoNS, not contaminant)** | | | 21,837,057 | 174,777,289 |
| CoNS spp. | 0/0 | 20/26 | | |
| **P073 (Blood culture grew *Streptococcus mitis* group)** | | | 14,867,751 | 74,667,179 |
| *Streptococcus mitis* group *spp.* | 0/0 | 0/0 | | |
| **P098 (Wound culture grew *Fusobacterium*, *Porphyromonas*; urine culture grew *A. urinae*)** | | | 19,727,410 | 172,614,577 |
| Fusobacterium spp. | 0/0 | 0/7 | | |
| *Porphyromonas* spp. | 2/6 | 12/57 | | |
| *Aerococcus urinae* | 0/0 | 0/2 | | |

| | | | | |
|---|---|---|---|---|
| **P152 (Blood culture grew CoNS, likely contaminant; urine culture grew *Klebsiella pneumoniae*)** | | | 17,866,699 | 103,288,138 |
| CoNS, likely contaminant | 1/1 | 0/1 | | |
| *Klebsiella pneumoniae* | 0/0 | 0/11 | | |
| **P118 (Bronchoalveolar culture grew *Staphylococcus aureus)*** | | | 30,271,177 | 262,289,640 |
| *Staphylococcus aureus* | 0/0 | 7/7 | | |

*Reads were processed using two alignment thresholds specified in Kraken: our original conservative threshold of 0.3, and a liberal threshold of 0.

CoNS = Coagulase Negative Staphylococcus

**Table S4.** Clinical relevance of all organisms found by mNGS that were not detected by hospital tests

| Patient | VirCapSeq-VERT and mNGS Organism* | Organism Clinically Relevant?† | | | mNGS oganism likely contaminant and/or misalignment?‡ | Standard-of-Care Microbiology, 0-5d After Presentation | Host Response | Final Diagnosis |
|---|---|---|---|---|---|---|---|---|
| | | R1 | R2 | R3 | | | | |
| P006 | *Escherichia coli* (37,26,12,Yes) | 5 | 5 | 5 | No | All negative. | Bacterial | Bacteremia – Source: Line |
| P013 | *Janthinobacterium* sp 1 2014MBL MicDiv (229,200,33,Yes) | 1 | 1 | 1 | Likely Contaminant and/or Misalignment | Blood Culture (D1): Escherichia coli (performed 2h after blood draw for study); Urine Culture (D1): 60,000 CFU/mL Streptococcus agalactiae (Group B). | Bacterial (Derivation Cohort)§ | Bacteremia – Source: Urine |

| | | | | | | | | |
|---|---|---|---|---|---|---|---|---|
| P020 | *Klebsiella pneumoniae* (85,69,4,Yes) | 3 | 5 | 5 | No | Blood Culture (D1): Escherichia coli; Abdominal Wound Culture (D1): 2+ Corynebacterium striatum | PCR Error | Bacteremia – Source: Unclear |
| | *Escherichia coli* (45,33,12,Yes) | In SOC Microbiology | | | Already identified by SOC microbiology. | | | |
| P021 | *Thermus scotoductus* (23,14,2,Yes) | 1 | 1 | 1 | Likely Contaminant and/or Misalignment | Blood Culture (D1): Streptococcus agalactiae (Group B); Urine Culture (D1): 30,000 CFU/mL Escherichia coli | Bacterial (Derivation Cohort)§ | Bacteremia - Unclear Etiology |
| P023 | Hepatitis C Virus (333,057/1,652) | 1 | 1 | 1 | N/A | All negative. | Noninfected (Derivation Cohort)§ | Malignancy - Metastatic Lung |
| | *Gardnerella vaginalis* (12,7,3,Yes) | 1 | 2 | 1 | No | | | |

| | | | | | | | | |
|---|---|---|---|---|---|---|---|---|
| P025 | Coxsackievirus B5 (30,324/181) | 3 | 5 | 5 | N/A | All negative. | Bacterial | Viral Syndrome |
| | *Corynebacterium maris* (27,18,5,Yes) | 3 | 1 | 2 | Likely Contaminant and/or Misalignment | All negative. | Bacterial | Viral Syndrome |
| P037 | *Prevotella denticola* (104,86,2,Yes) | 2 | 5 | 4 | No | Blood Culture (D1): Escherichia coli, Streptococcus anginosus group; Perianal Abscess Culture (D1): 3+ Streptococcus anginosus group; Blood Culture (D2): Streptococcus anginosus group; | Bacterial (Derivation Cohort)§ | Bacteremia – Source: Intra-Abdominal |
| | *Porphyromonas asaccharolytica* (73,58,2,Yes) | 2 | 5 | 4 | No | | | |
| | *Fusobacterium nucleatum* (61,47,3,Yes) | 3 | 5 | 4 | No | | | |
| | *Dialister pneumosintes* (8,3,0,No) | 2 | 2 | 2 | No | | | |

| | | | | | | | | |
|---|---|---|---|---|---|---|---|---|
| | | | | | | Perianal Abcsess Fluid Culture (D3): 2+ Bacteroides fragilis group; Perianal Abscess Fluid Culture (D3): 3+ Escherichia coli, 4+ Streptococcus anginosus group | | |
| P041 | *Fusobacterium nucleatum* (22,14,5,Yes) | 2 | 5 | 4 | No | All negative. | Bacterial | Intra-Abdominal Abscess |
| P057 | *Escherichia coli* (119,100,5,Yes) | 3 | 4 | 4 | No | Blood Culture (D1): Coagulase Negative Staphylococcus spp. | Noninfected (Derivation Cohort)§ | Allograft Rejection |

| | | | | | | | | |
|---|---|---|---|---|---|---|---|---|
| P058 | *Erwinia billingiae* (7,3,2,Yes) | 1 | 3 | 1 | Likely Contaminant and/or Misalignment | Urine Culture (D1): 30,000 CFU/mL Yeast | Bacterial | Post-Operative Fever vs. UTI |
| P061 | *Thermus scotoductus* (4,1,0,Yes) | 1 | 2 | 1 | Likely Contaminant and/or Misalignment | All negative. | Noninfected (Derivation Cohort)§ | Febrile Neutropenia - Unclear Etiology |
| P066 | *Propionibacterium* sp oral taxon 193 (8,4,2,Yes) | 1 | 1 | 1 | Likely Contaminant and/or Misalignment | Urine Culture (D1): >100,000 CFU/mL Escherichia coli | Bacterial (Derivation Cohort)§ | Pyelonephritis |
| | *Erwinia billingiae* (7,3,2,Yes) | 1 | 1 | 1 | Likely Contaminant and/or Misalignment | | | |
| P067 | *Propionibacterium acnes* (8,3,2,Yes) | 1 | 1 | 2 | Likely Contaminant and/or Misalignment | All negative. | Viral | Pneumonia |

| | | | | | | | | |
|---|---|---|---|---|---|---|---|---|
| P070 | *Staphylococcus warneri* (5440,5307,16,Yes) | 2 | 3 | 2 | No | All negative. | Noninfected (Derivation Cohort)§ | Pneumonia vs. Drug Reaction |
| | *Lactococcus lactis* (870,812,437,Yes) | 2 | 3 | 2 | No | | | |
| | *Actinomyces oris* (514,472,44,Yes) | 2 | 3 | 1 | No | | | |
| | *Streptococcus gordonii* (484,443,22,Yes) | 2 | 3 | 2 | No | | | |
| | *Rothia dentocariosa* (396,357,76,Yes) | 2 | 3 | 2 | No | | | |
| | *Veillonella parvula* (335,301,14,Yes) | 2 | 3 | 1 | No | | | |
| | *Streptococcus sanguinis* (265,237,12,Yes) | 2 | 3 | 2 | No | | | |
| | *Streptococcus mutans* (239,212,8,Yes) | 2 | 3 | 2 | No | | | |

| | | | | | |
|---|---|---|---|---|---|
| | *Streptococcus intermedius* (145,125,6,Yes) | 2 | 3 | 2 | No |
| | *Streptococcus oralis* (143,122,12,Yes) | 2 | 3 | 2 | No |
| | *Staphylococcus pasteuri* (119,99,7,Yes) | 2 | 3 | 2 | No |
| | *Fusobacterium nucleatum* (117,97,12,Yes) | 2 | 3 | 2 | No |
| | *Prevotella dentalis* (78,63,5,Yes) | 2 | 3 | 1 | No |
| | *Staphylococcus aureus* (53,39,10,Yes) | 2 | 3 | 3 | No |
| | *Selenomonas* sp oral taxon 920 (44,32,0,No) | 2 | 3 | 1 | No |
| | *Gardnerella vaginalis* (39,27,25,Yes) | 2 | 3 | 1 | No |

| | | | | | |
|---|---|---|---|---|---|
| | *Campylobacter gracilis* (38,27,5,Yes) | 2 | 3 | 2 | No |
| | Selenomonas sputigena (37,26,6,Yes) | 2 | 3 | 1 | No |
| | *Capnocytophaga* sp oral taxon 323 (32,22,5,Yes) | 2 | 3 | 2 | No |
| | *Leptotrichia* sp oral taxon 212 (22,14,5,Yes) | 2 | 3 | 1 | No |
| | *Tannerella* sp oral taxon HOT-286 (19,11,5,Yes) | 2 | 3 | 1 | No |
| | *Campylobacter concisus* (17,9,5,Yes) | 2 | 3 | 1 | No |
| | *Olsenella* sp oral taxon 807 (16,8,6,Yes) | 2 | 3 | 1 | No |
| | *Leptotrichia* sp oral taxon 847 (14,8,5,Yes) | 2 | 3 | 1 | No |

| | | | | | | | | |
|---|---|---|---|---|---|---|---|---|
| P071 | *Enterobacter cloacae* (11,6,2,Yes) | 1 | 2 | 3 | Possible Contaminant and/or Misalignment | All negative. | Noninfected | Coccidioides Meningitis |
| P073 | *Thermus scotoductus* (43,31,30,Yes) | 2 | 2 | 1 | Likely Contaminant and/or Misalignment | Blood Culture (D1): Streptococcus mitis group; Mouth Would Culture (D1): Acinetobacter baumannii; Blood Enzyme Immunoassay (D1): Aspergillus (Galactomannan) Antigen; Lesion PCR (D2): Herpes Simplex Virus 1 | Noninfected (Derivation Cohort)§ | Bacteremia – Source: Unclear |

| | | | | | | | | |
|---|---|---|---|---|---|---|---|---|
| P077 | *Thermus scotoductus* (11,5,0,Yes) | 1 | 1 | 1 | Likely Contaminant and/or Misalignment | Urine Culture (D5): >100,000 CFU/mL Escherichia coli | Bacterial (Derivation Cohort)§ | Necrotizing Pancreatitis |
| P083 | *Borrelia hermsii* (306,273,0,No) | 4 | 5 | 5 | No | Urine Culture (D1): >100,000 CFU/mL Coagulase Negative Staphylococcus spp. (not Staphylococcus saprophyticus) | Bacterial | Tick Borne Relapsing Fever |
| P084 | *Delftia acidovorans* (31,21,19,Yes) | 1 | 1 | 2 | Likely Contaminant and/or Misalignment | Abdominal Wound Culture (D1): 4+ Staphylococcus aureus | Noninfected (Derivation Cohort)§ | Intra-Abdominal Abscess |
| P086 | *Pseudomonas aeruginosa* (67,52,29,Yes) | 2 | 5 | 3 | No | Nasopharyngeal Swab PCR (D1): Rhinovirus. | Viral | Cystic Fibrosis Exacerbation |

| | | | | | | | | |
|---|---|---|---|---|---|---|---|---|
| P092 | *Streptococcus agalactiae* (21,13,0,No) | In SOC Microbiology | | | Already identified by SOC microbiology. | Blood Culture (D1): Streptococcus agalactiae (Group B), Escherichia coli; Urine Culture (D1): >100,000 CFU/mL Lactobacillus species, 20,000 CFU/mL Escherichia coli | Bacterial (Derivation Cohort)§ | Bacteremia – Source: Skin |
| | *Streptococcus anginosus* (9,4,2,Yes) | 4 | 5 | 5 | No | | | |
| P095 | *Propionibacterium acnes* (27,18,14,Yes) | 1 | 2 | 2 | Likely Contaminant and/or Misalignment | All negative. | Bacterial | Chemotherapy-Associated Fever |
| P101 | *Prevotella intermedia* (10,5,3,Yes) | 1 | 4 | 3 | Possible Contaminant and/or Misalignment | Blood Culture (D1): Viridans group Streptococci | Noninfected | Cholangitis |

| | | | | | | | | |
|---|---|---|---|---|---|---|---|---|
| P103 | *Enterobacter cloacae* (200,175,6,Yes) | In SOC Microbiology | | | Already identified by SOC microbiology. | Blood Culture (D1): Enterobacter cloacae complex | PCR Error | Bacteremia – Source: Unclear |
| | *Chroococcidiopsis thermalis* (51,38,3,Yes) | 2 | 3 | 1 | Likely Contaminant and/or Misalignment | | | |
| | *Enterobacter ludwigii* (15,8,0,No) | In SOC Microbiology | | | Already identified by SOC microbiology. | | | |
| P104 | *Pseudomonas* sp L1010 (120,100,5,Yes) | 1 | 2 | 2 | No | All negative. | Bacterial (Derivation Cohort)§ | Gastrostomy Tube Dysfunction |
| | *Pseudomonas fragi* (90,72,14,Yes) | 1 | 2 | 2 | No | | | |
| | *Acinetobacter baumannii* (72,56,31,Yes) | 1 | 2 | 2 | No | | | |

| | | | | | | | | |
|---|---|---|---|---|---|---|---|---|
| | *Leuconostoc citreum* (63,49,12,Yes) | 1 | 2 | 2 | No | | | |
| | *Psychrobacter alimentarius* (30,20,3,Yes) | 1 | 2 | 2 | No | | | |
| | *Cronobacter sakazakii* (18,11,0,No) | 1 | 2 | 2 | No | | | |
| | *Xanthomonas campestris* (17,10,3,Yes) | 1 | 2 | 1 | No | | | |
| P113 | *Klebsiella pneumoniae* (240,213,4,Yes) | 2 | 4 | 4 | No | All negative. | Bacterial (Derivation Cohort)§ | Ulcerative Colitis Flair |
| P115 | *Cupriavidus metallidurans* (5,2,0,Yes) | 1 | 3 | 2 | Likely Contaminant and/or Misalignment | All negative. | Bacterial (Derivation Cohort)§ | Malignancy - Leukemia |

| | | | | | | | | |
|---|---|---|---|---|---|---|---|---|
| P126 | *Morganella morganii* (29,20,2,Yes) | 4 | 5 | 5 | No | All negative. | Bacterial (Derivation Cohort)§ | Bacteremia – Source: Prostate |
| P133 | *Moraxella osloensis* (25,16,14,Yes) | 3 | 4 | 2 | Possible Contaminant and/or Misalignment | All negative. | Bacterial | Post-Operative Surgical Site Infection |
| P136 | *Kocuria palustris* (18,11,6,Yes) | 2 | 3 | 1 | No | All negative. | Noninfected | Febrile Neutropenia - Unclear Etiology |
| | *Brevibacterium linens* (11,6,2,Yes) | 2 | 3 | 1 | No | | | |
| | *Propionibacterium* sp oral taxon 193 (10,5,2,Yes) | 2 | 3 | 1 | No | | | |
| P137 | *Escherichia coli* (207,181,12,Yes) | In SOC Microbiology | | | Already identified by SOC microbiology. | Blood Culture (D1): Escherichia coli, Klebsiella oxytoca | Bacterial (Derivation Cohort)§ | Bacteremia – Source: Intra-Abdominal |

| | | | | | | | | |
|---|---|---|---|---|---|---|---|---|
| | *Clostridium perfringens* (101,83,2,Yes) | 4 | 5 | 5 | No | | | |
| P145 | *Enterobacter hormaechei* (123,103,2,Yes) | In SOC Microbiology | | | Already identified by SOC microbiology. | Blood Culture (D1): Enterobacter cloacae complex, Streptococcus anginosus group | Bacterial (Derivation Cohort)§ | Bacteremia – Source: Intra-Abdominal |
| | *Klebsiella pneumoniae* (26,17,4,Yes) | 4 | 4 | 5 | No | | | |
| | *Enterobacter cloacae* (16,9,2,Yes) | In SOC Microbiology | | | Already identified by SOC microbiology. | | | |
| | *Leclercia adecarboxylata* (8,4,2,Yes) | 2 | 2 | 2 | Possible Contaminant and/or Misalignment | | | |
| P153 | Hepatitis C Virus (423/2.6) | 1 | 1 | 3 | N/A | | | |

| | | | | | | | | |
|---|---|---|---|---|---|---|---|---|
| | *Klebsiella pneumoniae* (213,186,4,Yes) | In SOC Microbiology | | | Already identified by SOC microbiology. | Blood Culture (D1): Klebsiella pneumoniae; Serology (D2): Hepatitis B Surface Antibody | Bacterial (Derivation Cohort)§ | Bacteremia - Source Line |
| | *Lactococcus lactis* (57,44,8,Yes) | 1 | 1 | 3 | Possible Contaminant and/or Misalignment | | | |
| P154 | Streptococcus *mitis* (19,12,6,Yes) | 3 | 4 | 4 | No | Blood Culture (D1): Coagulase Negative Staphylococcus spp. | PCR Error | Febrile Neutropenia - Unclear Etiology |
| P163 | *Leptospira interrogans* (214,186,0,No) | 4 | 5 | 4 | No | All negative. | Bacterial | Leptospirosis |
| | *Xanthomonas campestris* (14,7,4,Yes) | 1 | 1 | 2 | Likely Contaminant and/or Misalignment | | | |

| | | | | | | | | |
|---|---|---|---|---|---|---|---|---|
| | *Anabaena* sp wa102 (13,7,3,Yes) | 1 | 1 | 2 | Likely Contaminant and/or Misalignment | | | |
| P166 | *Lactobacillus mucosae* (16,9,5,Yes) | 3 | 2 | 2 | No | All negative. | Bacterial | Balanitis |
| P171 | *Helicobacter pylori* (11,5,0,No) | 1 | 2 | 4 | No | All negative. | Bacterial | Cholangitis |
| P186 | *Escherichia* coli (*198*,171,141,Yes) | In SOC Microbiology | | | Already identified by SOC microbiology. | Blood Culture (D1): Escherichia coli; Urine Culture (D1): Escherichia coli | Bacterial (Derivation Cohort)§ | Bacteremia – Source: Urine |
| | *Pantoea* sp PSNIH1 (31,21,4,Yes) | 1 | 2 | 2 | Likely Contaminant and/or Misalignment | | | |
| P194 | *Haemophilus influenzae* (35,24,8,Yes) | 3 | 4 | 4 | No | All negative. | Bacterial | Pneumonia vs. Radiation Pneumonitis |

| | | | | | | | | |
|---|---|---|---|---|---|---|---|---|
| P197 | *Escherichia coli* (64, 49, 4, Yes) | 1 | 3 | 3 | No | All negative. | Noninfected (Derivation Cohort)§ | Abdominal Pain - Possible Calciphylaxis |
| | *Methyloversatilis* sp RAC08 (17, 10, 7, Yes) | 1 | 1 | 1 | Likely Contaminant and/or Misalignment | | | |

*mNGS (bacterial species) numbers represent (raw reads, estimated lower limit for the intensity of blood-associated reads, estimated upper limit for the intensity of contaminant reads, presence in negative controls). VirCapSeq-VERT (viruses) numbers represent (Raw Reads / Reads per 10,000 Host Subtracted Reads).

†Indicates whether physicians classified organism as clinically relevant to the patient's presentation while blinded to host response results. 1 = No, 2 = Probably No, 3 = Unsure, 4 = Probably Yes, 5 = Yes. R= Reviewer.

‡A fourth unblinded physician assessed whether mNGS organisms were likely to be contaminants and/or misalignments after the completion of our main chart review. Answer choices included No, Possible Contaminant and/or Misalignment, and Likely Contaminant and/or Misalignment. This physician considered medical charts, mNGS results, VirCapSeq-VERT results, host response results, and all classifications made by the three physician chart reviewers in the main chart review.

§Indicates patients from derivation cohort who had their host response results used to re-establish cutoffs for host response scores (see Fig. S4B).

**Table S5.** Clinical relevance of all viruses found by VirCapSeq-VERT that were not detected by hospital tests: Summary Table

| **VirCapSeq-VERT Organisms Not Detected by Standard-of-Care Microbiology*** | **Likely Clinically Relevant†** | **Uncertain Clinical Relevance** | **Likely Viral Reactivation and/or Chronic Infection, Not Clinically Relevant** |
|---|---|---|---|
| Human herpesvirus 6 | 0 | 2 | 2 |
| Epstein-Barr virus‡ | 1 | 0 | 7 |
| Hepatitis C virus | 0 | 2 | 6 |
| Hepatitis B virus | 0 | 0 | 1 |
| BK virus‡ | 0 | 0 | 2 |
| Trichodysplasia spinulosa-associated polyomavirus | 0 | 0 | 1 |
| Coxsackievirus B4 | 0 | 1 | 0 |
| Coxsackievirus B5 | 1 | 0 | 0 |
| Coxsackievirus A6 | 1 | 0 | 0 |
| Human parvovirus B19 | 0 | 1 | 0 |

*Includes tests performed within 5 days after presentation.

**†**Patients in the “Likely Clinically Relevant” column had VirCapSeq-VERT organisms which were classified as clinically relevant or probably clinically relevant to the patient’s presentation by physician consensus while blinded to host response results. Patients were classified into the remaining two columns by a fourth unblinded physician after the completion of our main chart review. This physician considered medical charts, mNGS results, VirCapSeq-VERT results, host response results, and all classifications made by the three physician chart reviewers in the main chart review.

‡One patient had both viral reactivation with Epstein-Barr virus and possible chronic infection with BK virus. There were 27 patients total.

Clinical details for all patients in this Table are presented in Table S6.

**Table S6.** Clinical relevance of all viruses found by VirCapSeq-VERT that were not detected by hospital tests: Clinical details

| Patient | VirCapSeq-VERT and mNGS Organism* | Organism Clinically Relevant?† | | | Comments on Clinical Relevance‡ | Pos. Standard-of-Care Microbiology, 0-5d After Presentation | Host Response | Final Diagnosis |
|---|---|---|---|---|---|---|---|---|
| | | **R1** | **R2** | **R3** | | | | |
| P003 | Coxsackievirus A6 (33,309,209/8,185) | 3 | 4 | 5 | Likely Clinically Relevant | All negative. | Bacterial | Viral Syndrome |
| P004 | Human herpesvirus 6 (4,519/9.7) | 2 | 4 | 3 | Likely Viral Reactivation and/or Chronic Infection, Not Clinically Relevant | All negative. | PCR Error | Febrile Neutropenia Unclear Etiology |
| P015 | Human herpesvirus 6 (3,550/20.5) | 1 | 4 | 2 | Likely Viral Reactivation and/or Chronic Infection, Not Clinically Relevant | Serology (D4): Hepatitis A virus IgG | Bacterial | Febrile Neutropenia Relapsed A[illegible] |

| | | | | | | | | |
|---|---|---|---|---|---|---|---|---|
| P019 | Hepatitis C virus (2,785,501 / 3,423.27) | 1 | 2 | 2 | Likely Viral Reactivation and/or Chronic Infection, Not Clinically Relevant | Urine Culture (D1, D2): >100,000 CFU/mL *Citrobacter freundii* complex, >100,000 CFU/mL *Enterococcus* species | Bacterial (Derivation Cohort)§ | Pyelonephrit |
| | *Citrobacter freundii* (115,97,7,Yes) | In SOC Microbiology | | | N/A | | | |
| P023 | Hepatitis C virus (333,057/1,652) | 1 | 1 | 1 | Likely Viral Reactivation and/or Chronic Infection, Not Clinically Relevant | All negative. | Noninfected (Derivation Cohort)§ | Malignancy - Metastatic Lung |
| | *Gardnerella vaginalis* (12,7,3,Yes) | 1 | 2 | 1 | N/A | | | |
| P025 | Coxsackievirus B5 (30,324/181) | 3 | 5 | 5 | Likely Clinically Relevant | All negative. | Bacterial | Viral Syndrome |
| | *Corynebacterium maris* (27,18,5,Yes) | 3 | 1 | 2 | N/A | | | |

| | | | | | | | | |
|---|---|---|---|---|---|---|---|---|
| P039 | Epstein-Barr virus (239,712/1,338.8) | 2 | 1 | 3 | Likely Viral Reactivation and/or Chronic Infection, Not Clinically Relevant | All negative. | Bacterial | Malignancy - Lymphoma |
| P040 | Epstein-Barr virus (336 / 1.46) | 1 | 2 | 2 | Likely Viral Reactivation and/or Chronic Infection, Not Clinically Relevant | Plasma PCR (D1, D4): Cytomegalovirus; Serology (D1): Cytomegalovirus IgG, Cytomegalovirus IgM, Epstein-Barr virus VCA IgG, Epstein-Barr virus EBNA IgG; Serology (D3): *Coxiella burnetti* (Q Fever IGG Phase). | Bacterial (Derivation Cohort)§ | Infectious Mononucleo |
| | Cytomegalovirus (2,936/12.8) | In SOC Microbiology | | | N/A | | | |
| | *Cupriavidus metallidurans* (8,3,1,Yes) | 1 | 1 | 2 | N/A | | | |
| P052 | Epstein-Barr virus (1,673/6.9) | 1 | 2 | 2 | Likely Viral Reactivation and/or Chronic Infection, Not Clinically Relevant | Blood Culture (*D1): Streptococcus anginosus,* | Bacterial (Derivation Cohort)§ | Cholecysitits Hepatic Abscess |

| | | | | | | | | |
|---|---|---|---|---|---|---|---|---|
| | BK Virus (2,384/9.9) | 1 | 2 | 1 | Likely Viral Reactivation and/or Chronic Infection, Not Clinically Relevant | *Haemophilus influenzae*; Bile Fluid Culture (D4): 4+ *Enterococcus faecalis*, 2+ *Streptococcus mitis* | | |
| | *Haemophilus influenzae* (190,165,2,Yes) | In SOC Microbiology | | | N/A | | | |
| P076 | Epstein-Barr virus (11,422/140) | 2 | 4 | 4 | Likely clinically Relevant. (However, possibility for viral reactivation remains.) | All negative. Note: The following test was also identified: Urine Culture (D -3): > 100,000 CFU/mL Enterococcus Species | Bacterial (Derivation Cohort)§ | UTI vs. Malignancy - Lymphoma |
| P081 | Hepatitis C virus (88,841/389) | 1 | 1 | 1 | Likely Viral Reactivation and/or Chronic Infection, Not Clinically Relevant | Urine Culture (D1): >100,000 CFU/mL Escherichia coli | PCR Error | Pyelonephrit |

| | | | | | | | | |
|---|---|---|---|---|---|---|---|---|
| P089 | BK virus (541/1.1) | 1 | 1 | 1 | Likely Viral Reactivation and/or Chronic Infection, Not Clinically Relevant | Blood Culture (D1): *Escherichia coli*; Urine Culture (D1): 50,000 CFU/mL *Escherichia coli*. | Bacterial | Bacteremia - Source: Urin |
| P096 | Coxsackievirus B4 (3,097/20) | 2 | 3 | 3 | Uncertain: Presentation fits with UTI | Urine Culture (D1): 100,000 CFU/mL *Escherichia coli* | Bacterial (Derivation Cohort)§ | Pyelonephri |
| P107 | Human parvovirus B19 (378/0.4) | 1 | 2 | 2 | Uncertain: Presentation fits with Norovirus infection. However, patient exposed to young children. | Stool PCR (D1): Norovirus | Bacterial (Derivation Cohort)§ | Diarrhea - Infectious |
| P112 | Epstein-Barr virus (91/1.9) | 1 | 3 | 1 | Likely Viral Reactivation and/or Chronic Infection, Not Clinically Relevant | Urine Culture (D4): >100,000 CFU/mL *Escherichia coli* | Bacterial | Pyelonephri |

| | | | | | | | | |
|---|---|---|---|---|---|---|---|---|
| P114 | Hepatitis C virus (34,254/241) | 1 | 2 | 1 | Likely Viral Reactivation and/or Chronic Infection, Not Clinically Relevant | Blood Culture (D1): Probable Coagulase Negative *Staphylococcus* spp. | Bacterial (Derivation Cohort)§ | Malignancy - Metastatic Prostate |
| P130 | Hepatitis C virus (3,584,985/2,600) | 1 | 1 | 2 | Likely Viral Reactivation and/or Chronic Infection, Not Clinically Relevant. Documented HCV chronic infection. | Blood Culture (D1): Coagulase Negative *Staphylococcus* spp. | Bacterial | Allograft Rejection |
| P146 | Trichodysplasia spinulosa-associated polyomavirus (698/2.8) | 1 | 1 | 2 | Likely Viral Reactivation and/or Chronic Infection. Possibly due to steroids and TNF-alpha suppression. No documented skin lesions. | All negative. | Bacterial (Derivation Cohort)§ | Crohn's Flai vs. Small Bowel Obstruction |
| P148 | Epstein-Barr virus (275/1.6) | 1 | 3 | 2 | Likely Viral Reactivation and/or Chronic Infection, Not Clinically Relevant | Blood Culture (D2, D4): *Pseudomonas aeruginosa*; Respiratory | Bacterial (Derivation Cohort)§ | Bacteremia - Source: Respiratory |

| | | | | | | | | |
|---|---|---|---|---|---|---|---|---|
| | *Pseudomonas aeruginosa* (2067, 1979, 46, Yes) | In SOC Microbiology | | | N/A | Culture (D1): *Pseudomonas aeruginosa.* | | |
| P149 | Hepatitis C virus (14,080,841/7,060.5) | 1 | 1 | 2 | Likely Viral Reactivation and/or Chronic Infection, Not Clinically Relevant | Nasopharyngeal Swab PCR (D1): Influenza A 2009 H1N1 | Viral (Derivation Cohort)§ | URI |
| P150 | Hepatitis B virus (49,462,447 / 9,210.54) | 1 | 1 | 2 | Likely Viral Reactivation and/or Chronic Infection, Not Clinically Relevant | Urine Culture (D1): 50,000 CFU/mL *Escherichia coli* | Bacterial (Derivation Cohort)§ | Pyelonephrit |
| P152 | Epstein-Barr virus (119 / 1.00) | 1 | 2 | 2 | Likely Viral Reactivation and/or Chronic Infection, Not Clinically Relevant | Blood Culture (D1): Coagulase Negative *Staphylococcus* spp.; Urine Culture (D1): >100,000 CFU/mL *Klebsiella pneumoniae* | Bacterial (Derivation Cohort)§ | Pyelonephrit |

| | | | | | | | | |
|---|---|---|---|---|---|---|---|---|
| P153 | Hepatitis C virus (423 / 2.57) | 1 | 1 | 3 | Uncertain: Clinical history does not align with chronic HCV infection. | Blood Culture (D1): *Klebsiella pneumoniae;* Serology (D2): Hepatitis B Surface Antibody | Bacterial (Derivation Cohort)§ | Bacteremia [illegible] Source: Line[illegible] |
| | *Klebsiella pneumoniae* (213, 186, 4, Yes) | In SOC Microbiology | | | N/A | | | |
| | Lactococcus lactis (57, 44, 8, Yes) | 1 | 1 | 3 | N/A | | | |
| P156 | Epstein-Barr virus (1811/2.7) | In SOC Microbiology | | | Likely Viral Reactivation | Serology (D1): Epstein-Barr virus Monospot Antibody Test; Plasma PCR (D2): Cytomegalovirus, Epstein-Barr virus | Bacterial | Post-Operat[illegible] Fever |
| | Human herpesvirus 6 (21,003/30.8) | 2 | 4 | 2 | Uncertain: Clinically Relevant or Viral Reactivation | Serology (D1): Epstein-Barr virus Monospot | Bacterial | Post-Operat[illegible] Fever |

| | | | | | | | | |
|---|---|---|---|---|---|---|---|---|
| | | | | | | Antibody Test; Plasma PCR (D2): Cytomegalovirus, Epstein-Barr virus | | |
| P164 | Hepatitis C virus (6,097/22.5) | 1 | 2 | 2 | Uncertain: Clinical history does not align with chronic HCV infection. | Plasma PCR (D1): Cytomegalovirus | Bacterial (Derivation Cohort)§ | Infectious Mononucleo[illegible] |
| | Cytomegalovirus (4,502/16.6) | In SOC Microbiology | | | Already identified by SOC microbiology. | Plasma PCR (D1): Cytomegalovirus | Bacterial (Derivation Cohort)§ | Infectious Mononucleo[illegible] |
| P165 | Human herpesvirus 6 (288 / 1.42) | 2 | 4 | 2 | Uncertain: Clinically Relevant or Viral Reactivation | All negative. | Bacterial | Pneumonia [illegible] Drug Reacti[illegible] |
| P187 | Hepatitis C virus (27,407,996 / 3,987.89) | 1 | 1 | 2 | Likely Viral Reactivation and/or Chronic Infection, Not Clinically Relevant | Serology (D1): Hepatitis B Surface Antigen | Viral (Derivation Cohort)§ | Pneumonia [illegible] Aspiration Pneumonitis |

*mNGS (bacterial species) numbers represent (raw reads, estimated lower limit for the intensity of blood-associated reads, estimated upper limit for the intensity of contaminant reads, presence in negative controls). VirCapSeq-VERT (viruses) numbers represent (Raw Reads / Reads per 10,000 Host Subtracted Reads).

†Indicates whether physicians classified virus as clinically relevant to the patient's presentation while blinded to host response results. 1 = No, 2 = Probably No, 3 = Unsure, 4 = Probably Yes, 5 = Yes. R= Reviewer.

‡Comments were made by a fourth unblinded physician as to whether viruses were clinically relevant, chronic infections and/or viral reactivations that were not clinically relevant, or of uncertain clinical relevance. This physician considered medical charts, mNGS results, VirCapSeq-VERT results, host response results, and all classifications made by the three physician chart reviewers in the main chart review.

§Indicates patients from derivation cohort who had their host response results used to re-establish cutoffs for host response scores (see Fig. S4B).

**Table S7. Clinical details for four cases in which mNGS and VirCapSeq-VERT had the greatest influence in changing physician classifications**

<table>
<tr><th>Patient*</th><th>Presentation</th><th>SOC Micro-biology</th><th>Anti-biotics?</th><th>mNGS and VirCapSeq-VERT</th><th>Host Response</th><th>Chart Review Classifications</th></tr>
<tr><td>P003</td><td>Male with fever, myalgia, and erythematous lesions suspected from bug bites. Presumed cellulitis. Discharged home after IV fluids, no follow-up data.</td><td>Blood culture neg.</td><td>Yes</td><td>mNGS: Negative<br>VirCapSeq-VERT: Coxsackievirus A6</td><td>Bacterial</td><td>
<table>
<tr><th></th><th colspan="3">Rev. 1</th><th colspan="3">Rev. 2</th><th colspan="3">Rev. 3</th></tr>
<tr><th></th><th>PI</th><th>PII</th><th>PIII</th><th>PI</th><th>PII</th><th>PIII</th><th>PI</th><th>PII</th><th>PIII</th></tr>
<tr><td>Infected?</td><td>3</td><td>3</td><td>3</td><td>3</td><td>4</td><td>4</td><td>4</td><td>5</td><td>5</td></tr>
<tr><td>Bacterial</td><td>3</td><td>3</td><td>3</td><td>2</td><td>2</td><td>3</td><td>4</td><td>1</td><td>2</td></tr>
<tr><td>Viral?</td><td>3</td><td>3</td><td>3</td><td>3</td><td>4</td><td>4</td><td>2</td><td>5</td><td>5</td></tr>
</table>
</td></tr>
<tr><td>P025</td><td>Female with fever, vomiting, diarrhea, and headache. Presumed UTI and/or viral syndrome. Discharged home after IV fluids, no follow-up data.</td><td>Blood and urine cultures neg. Pyuria on urinalysis.</td><td>Yes</td><td>mNGS: Negative<br>VirCapSeq-VERT: Coxsackievirus B5</td><td>Bacterial</td><td>
<table>
<tr><th></th><th colspan="3">Rev. 1</th><th colspan="3">Rev. 2</th><th colspan="3">Rev. 3</th></tr>
<tr><th></th><th>PI</th><th>PII</th><th>PIII</th><th>PI</th><th>PII</th><th>PIII</th><th>PI</th><th>PII</th><th>PIII</th></tr>
<tr><td>Infected?</td><td>2</td><td>3</td><td>3</td><td>4</td><td>5</td><td>5</td><td>5</td><td>5</td><td>5</td></tr>
<tr><td>Bacterial</td><td>2</td><td>3</td><td>3</td><td>2</td><td>1</td><td>1</td><td>4</td><td>2</td><td>2</td></tr>
<tr><td>Viral?</td><td>1</td><td>3</td><td>3</td><td>4</td><td>5</td><td>5</td><td>2</td><td>5</td><td>5</td></tr>
</table>
</td></tr>
</table>

<table>
<tr>
<td>P083</td>
<td>Male with fever, dry cough, and BPH-related urinary retention returning from field work in a mountainous region. Presumed urosepsis. Discharged home; improved on 3-day follow-up.</td>
<td>Blood and urine cultures neg. Pyuria on urinalysis.</td>
<td>Yes</td>
<td>mNGS: Borrelia hermsii<br><br>VirCapSeq-VERT: Negative</td>
<td>Bacterial</td>
<td>
<table>
<tr><th></th><th colspan="3">Rev. 1</th><th colspan="3">Rev. 2</th><th colspan="3">Rev. 3</th></tr>
<tr><th></th><th>PI</th><th>PII</th><th>PIII</th><th>PI</th><th>PII</th><th>PIII</th><th>PI</th><th>PII</th><th>PIII</th></tr>
<tr><td>Infected?</td><td>4</td><td>5</td><td>5</td><td>5</td><td>5</td><td>5</td><td>4</td><td>5</td><td>5</td></tr>
<tr><td>Bacterial</td><td>4</td><td>5</td><td>5</td><td>3</td><td>5</td><td>5</td><td>4</td><td>5</td><td>5</td></tr>
<tr><td>Viral?</td><td>2</td><td>1</td><td>1</td><td>3</td><td>1</td><td>1</td><td>1</td><td>1</td><td>1</td></tr>
</table>
</td>
</tr>
<tr>
<td>P163</td>
<td>Male with fever, headache, and diarrhea upon return from Sri Lanka. Presumed viral infection. Discharged home under strict return precautions; symptoms continued but improved on 8-day follow-up.</td>
<td>Blood culture, dengue, and malaria tests neg.</td>
<td>No</td>
<td>mNGS: Leptospira interrogans<br><br>VirCapSeq-VERT: Negative</td>
<td>Bacterial</td>
<td>
<table>
<tr><th></th><th colspan="3">Rev. 1</th><th colspan="3">Rev. 2</th><th colspan="3">Rev. 3</th></tr>
<tr><th></th><th>PI</th><th>PII</th><th>PIII</th><th>PI</th><th>PII</th><th>PIII</th><th>PI</th><th>PII</th><th>PIII</th></tr>
<tr><td>Infected?</td><td>4</td><td>5</td><td>5</td><td>5</td><td>5</td><td>5</td><td>5</td><td>5</td><td>5</td></tr>
<tr><td>Bacterial</td><td>2</td><td>4</td><td>4</td><td>2</td><td>5</td><td>5</td><td>2</td><td>4</td><td>5</td></tr>
<tr><td>Viral?</td><td>4</td><td>2</td><td>2</td><td>4</td><td>1</td><td>1</td><td>4</td><td>2</td><td>1</td></tr>
</table>
</td>
</tr>
</table>

<table>
<tr><td rowspan="2">Legend</td><th>Phase I (PI)</th><th>Phase II (PII)</th><th>Phase III (PIII)</th><th>No</th><th>Probably No</th><th>Unsure</th><th>Probably Yes</th><th>Yes</th></tr>
<tr><td>Medical Charts Only</td><td>+mNGS, VirCapSeq-VERT</td><td>+Host Response</td><td>1</td><td>2</td><td>3</td><td>4</td><td>5</td></tr>
</table>

*Patients were identified by the following criteria: 1) An organism was revealed by mNGS or VirCapSeq-VERT that was not previously detected by standard-of-care microbiology, and 2) physicians classified the organism as clinically relevant or probably clinically relevant by consensus, and 3) mNGS or VirCapSeq-VERT results led at least one physician to increase their clinical suspicion for bacterial or viral infection by at least two points on a five-point scale to “Yes.”

SOC = Standard-of-care; mNGS = metagenomic next generation sequencing; BPH = benign prostate hypertrophy, Rev = physician chart reviewer

**Table S8.** Physician interpretation and host response results for patients originally classified as noninfected or probably noninfected, and found by mNGS to have bacterial sequences in plasma

| Patient* | mNGS Organisms‡ | mNGS Organism Clinically Relevant? ¶ | | | Clinical Comments | Antibiotics\\ | Improved?** | Host Response |
|---|---|---|---|---|---|---|---|---|
| | | R1 | R2 | R3 | | | | |
| P023 | *Gardnerella. vaginalis* (12, 7, 3) | 1 | 2 | 1 | Female with COPD exacerbation, lymphangitic carcinomatosis. | N | Y | Not Infected (Derivation Cohort)†† |
| P057 | *Escherichia coli* (119, 100, 5) | 3 | 4 | 4 | Renal transplant rejection. | N | Y | Not Infected (Derivation Cohort)†† |
| P070 | *Staphylococcus warneri* (5440, 5307, 16) | 2 | 3 | 2 | Likely Nivolumab-associated pneumonitis. +Hemoptysis, severe gingivitis. | Y | Y | Not Infected (Derivation Cohort)†† |
| | *Lactococcus lactis* (870, 812, 437) | 2 | 3 | 2 | | | | |
| | *Actinomyces oris* (514, 472, 44) | 2 | 3 | 1 | | | | |
| | *Streptococcus gordonii (484, 443, 22)* | 2 | 3 | 2 | | | | |

| | | | | | | | | |
|---|---|---|---|---|---|---|---|---|
| | *Rothia dentocariosa* (396, 357, 76) | 2 | 3 | 2 | | | | |
| | +19 Additional Organisms§ | | | | | | | |
| P104 | *Pseudomonas* sp. L1010 (120, 100, 5) | 1 | 2 | 2 | Clogged gastronomy tube, extensive bowel resection. +Total parenteral nutrition. | N | Y | Bacterial (Derivation Cohort)†† |
| | *Pseudomonas fragi* (90, 72, 14) | 1 | 2 | 2 | | | | |
| | *Acinetobacter baumannii* (72, 56, 31) | 1 | 2 | 2 | | | | |
| | *Leuconostoc citreum* (63, 49, 12) | 1 | 2 | 2 | | | | |
| | *Psychrobacter alimentarius* (30, 20, 3) | 1 | 2 | 2 | | | | |
| | *Cronobacter sakazakii* (18, 11, 0) | 1 | 2 | 2 | | | | |
| | *Xanthomonas campestris* (17, 10, 3) | 1 | 2 | 1 | | | | |
| P113 | *Klebsiella pneumoniae* (240, 213, 4) | 2 | 4 | 4 | Ulcerative Colitis flare. +Recent *C. difficiles* colitis on Vancomycin taper. | N | Y | Bacterial (Derivation Cohort)†† |
| P136 | *Kocuria palustris* (18, 11, 6) | 2 | 3 | 1 | | Y | Y | Not Infected |

| | | | | | | | | |
|---|---|---|---|---|---|---|---|---|
| | *Brevibacterium linens* (11, 6, 2) | 2 | 3 | 1 | Neutropenic fever following chemotherapy. +Mucositis, oral ulcers. | | | |
| | *Propionibacterium* sp. oral taxon 193 (10, 5, 2) | 2 | 3 | 1 | | | | |
| P194 | *Haemophilus influenzae* (35, 24, 8) | 3 | 4 | 4 | Pneumonia or radiation pneumonitis | N | Y | Bacterial |
| P197 | *Escherichia coli* (64, 49, 4) | 1 | 3 | 3 | Abdominal pain and chronic wounds likely due to calciphylaxis. | N | Y | Not Infected (Derivation Cohort)†† |
| | *Methyloversatilis* sp. RAC08 (17, 10, 7)[b] | 1 | 1 | 1 | | | | |

| No | Probably No | Unsure | Probably Yes | Yes |
|---|---|---|---|---|
| 1 | 2 | 3 | 4 | 5 |

*Patients were identified by the following criteria: 1) patient had consensus classification as either noninfected or probably noninfected by physicians while blinded to mNGS, VirCapSeq-VERT, and host response data; and 2) the patient had a positive mNGS result for an organism not detected by standard-of-care microbiology within 5 days after presentation.

‡The three numbers in parentheses indicate (raw reads, estimated lower limit for the intensity of blood-associated reads, estimated upper limit for the intensity of contaminant reads). All patients in this table had negative VirCapSeq-VERT results,

except for Pt_023, who had a positive result for hepatitis C Virus. Additionally, all patients did not have any positive, clinically relevant standard-of-care microbiology results.

§Patient P070 had 19 additional oral-related organisms identified by mNGS. Full results are provided in supplementary attachment 1.

¶Clinical relevance determined by three physician chart reviewers (R1-3) who examined mNGS and VirCapSeq-VERT results in the context of the entire medical record, while blinded to host response results.

\\Indicates whether antibiotics were prescribed to the patient for the sepsis-like illness that prompted their ED visit and/or admission, as documented by medical chart data.

**Indicates whether the patient improved, as documented by medical chart data.

††Indicates patients from derivation cohort who had their host response results used to re-establish cutoffs for host response scores (see Fig. S4B).

COPD = Chronic obstructive pulmonary disease.
